# Maternal repression of imprinted genes in endosperm by gene body demethylation

**DOI:** 10.64898/2026.09.29.755427

**Authors:** Chiragh Dewan, Yirui Sun, Jonathan I. Gent

## Abstract

Removal of TE-like DNA methylation from regulatory elements drives canonical genomic imprinting of both maternally and paternally expressed genes in endosperm (MEGs and PEGs). In both cases, the maternal copies are demethylated, allowing binding of either activating or repressing factors. Here we report that removal of CG-only gene body methylation also drives imprinting, resulting in maternally repressed PEGs as the most common form of imprinting in maize. These body methylated PEGs express broadly across tissues and lack H3K27me3 except in endosperm, where maternal copies acquire it in domains extending through and beyond genes. We also found that partial gene body demethylation is widespread in endosperm, usually does not result in imprinted expression, and is mediated by DNA glycosylases/lyases (DNGs). These and other data indicate that gene body methylation can shield select genes from PRC2 repression, and its maternal removal by DNGs explains the majority of PEGs in both maize and rice but has a weak impact in *Arabidopsis thaliana*. We hypothesize that increased gene expression noise arising from gene body demethylation–both with and without imprinting–is beneficial in endosperm.

## INTRODUCTION

Epigenetic differences in male and female gametes can lead to differential maternal and paternal gene expression after fertilization. In an extreme form, genomic imprinting happens at the level of whole chromosomes, with examples from diverse organisms including insects, mammals, and liverworts [1–3]. More widespread genomic imprinting happens at the level of individual genes, as first rigorously demonstrated by imprinting of *r1*, a maize endosperm pigmentation gene that is preferentially maternally expressed [4,5]. Evidence for paternal-preferred imprinting came from demonstrations that specific segments of paternally inherited chromosomes are required for maize endosperm development [6,7]. Subsequently, another example of a maternally preferred locus, *dzr1*, was discovered in maize [8], as were multiple loci in mice, e.g., *Tme* [9]. DNA methylation patterns associated with imprinting in mice pointed toward a role for DNA methylation and demethylation, which was confirmed by loss of imprinting in a methylation defective mouse mutant [10]. At about the same time, imprinting of a family of protein storage genes in maize endosperm was linked to maternal demethylation [11]. A major discovery soon followed from work in the model species *Arabidopsis thaliana* (hereafter arabidopsis): identification of a DNA glycosylase/lyase (DNG) responsible for maternal DNA demethylation in imprinting [12,13]. The gene behind a maize imprinting mutant that Kermicle originally characterized (*maternal derepression of r1*, or *mdr1*) was later discovered to also encode a DNG [4,14,15].

Since methylation of cis-regulatory elements can inhibit both transcriptional activators and repressors, demethylation can lead to either activation or repression, thus producing maternally and paternally expressed genes (MEGs and PEGs). Histone H3 lysine 27 trimethylation (H3K27me3), established and maintained by Polycomb Repressive Complex 2 (PRC2), provides a mechanism for maternal repression of PEGs, not only in canonical imprinting by maternal demethylation, but also in other forms of imprinting that do not involve maternal demethylation [16]. In diverse plants and animals, coding DNA of a subset of genes is marked by DNA methylation specifically in the CG context (sometimes referred to as CpG to distinguish dinucleotides from C-G base pairing). This CG-only form of methylation, called gene body methylation (gbM), is distinct from TE-like DNA methylation (teM) in gene bodies, which occurs in CG context together with CHG and to a lesser extent CHH as well (where H = A, T, or C) [17,18]. teM is linked to other constitutive heterochromatin modifications and is often an indicator of dubious gene annotations, but it can also be a signature of nearly constitutive gene silencing but high and specific expression in either pollen or endosperm in maize [18–20]. When teM extends upstream of gene bodies into promoters, its maternal demethylation produces teM-MEGs [20].

Due to a lack of functional evidence for specific methyl binding proteins that recognize gbM, widespread indirect effects of methylation mutants on transcription, difficulty in separating methylation changes from linked genetic changes, and the observation that some closely related plants lack it altogether, it is not clear what consequence gbM has on transcription [21–24]. As in other plants, in maize it is a strong indicator of broad and stable gene expression [18]. Given the role of H3K27me3 in tissue specific repression, one would expect it to be absent from constitutively expressed gbM genes [25], but evidence from both plants and animals suggests a more direct antagonism between gbM and H3K27me3. In particular, H3K27me3 is recruited to some newly CG hypomethylated regions [26–29]. Consistent with this, a connection between maternal CG demethylation of gene bodies and increased H3K27me3 has been observed in PEGs in both rice and maize endosperm [30–32].

These findings raise the question of whether active demethylation of gene bodies promotes PRC2 or whether PRC2 inhibits gene body methylation in endosperm. The latter would be more consistent with canonical mechanisms of imprinting determined by epigenetic features of cis-regulatory elements. In either case, it raises the additional questions of whether gene body methylation itself is important for imprinting or a side effect of demethylation of cis regulatory elements in select genes. To answer these questions, we identified high-confidence imprinted genes from recently released endosperm RNA-seq data from bidirectional crosses between B73 and eight inbred maize lines and compared imprinting status with genic DNA methylation. We found that maternal gene body demethylation is not limited to PEGs but is a general feature of endosperm, usually partial demethylation and without detectable effects on gene expression. In particular, maternal CG methylation was reduced 24% relative to paternal methylation on average in biparentally expressed (non-imprinted) gbM genes. Mutant analysis indicated that DNGs mediate maternal gene body demethylation of both biparentally expressed genes and PEGs, but maternal H3K27me3 was only detectable in PEGs. Taken together with other data on PRC2 in imprinting, these results indicate that gbM protects paternal copies of select genes from PRC2 repression in endosperm while maternal demethylation exposes them. This phenomenon accounts for approximately half of strongly imprinted genes in rice (150 gbM PEGs of 313 identified imprinted genes) and maize (102 of 180), but it is rare or negligible in arabidopsis (11 of 106).

## RESULTS

### Identification of imprinted genes in maize

To produce a high confidence list of imprinted genes, we used the most extensive dataset of hybrid endosperm mRNA-seq available [33]. This data were produced using genetic polymorphisms between parental genomes to quantify parent-specific gene expression in endosperm from bidirectional crosses between B73 and eight other inbred stocks with high-quality genome assemblies and gene annotations [34,35]. The reported expression measurements for each homologous parental gene (syntelog) were produced by mapping mRNA-seq reads to concatenated reference genomes for each pair of genomes and counting reads for each. Given the two-to-one maternal-to-paternal karyotype of endosperm, unbiased biparental expression should yield a maternal ratio of 2/3. We defined MEGs as having a threefold or greater than expected maternal expression, corresponding to a maternal ratio of 6/7. Similarly, we defined PEGs as having threefold or greater paternal expression than expected, corresponding to a maternal ratio of 2/5. We defined genes with intermediate values as biparental. To account for the complication that tandemly duplicated genes with different numbers of copies in different genomes can create multiple potential syntelog pairs, we required that genes with multiple syntelogs had the same imprinting call for each of their potential syntelog pairs or were left uncategorized. To account for both developmental staging variation and syntelog expression differences, we included only genes with reproducible expression patterns in at least two hybrids and without conflicting expression patterns in any hybrids. This method yielded 27 MEGs, 153 PEGs, and 11,331 biparentals (Fig. 1a and Supplemental Dataset 1).

**Figure 1:**
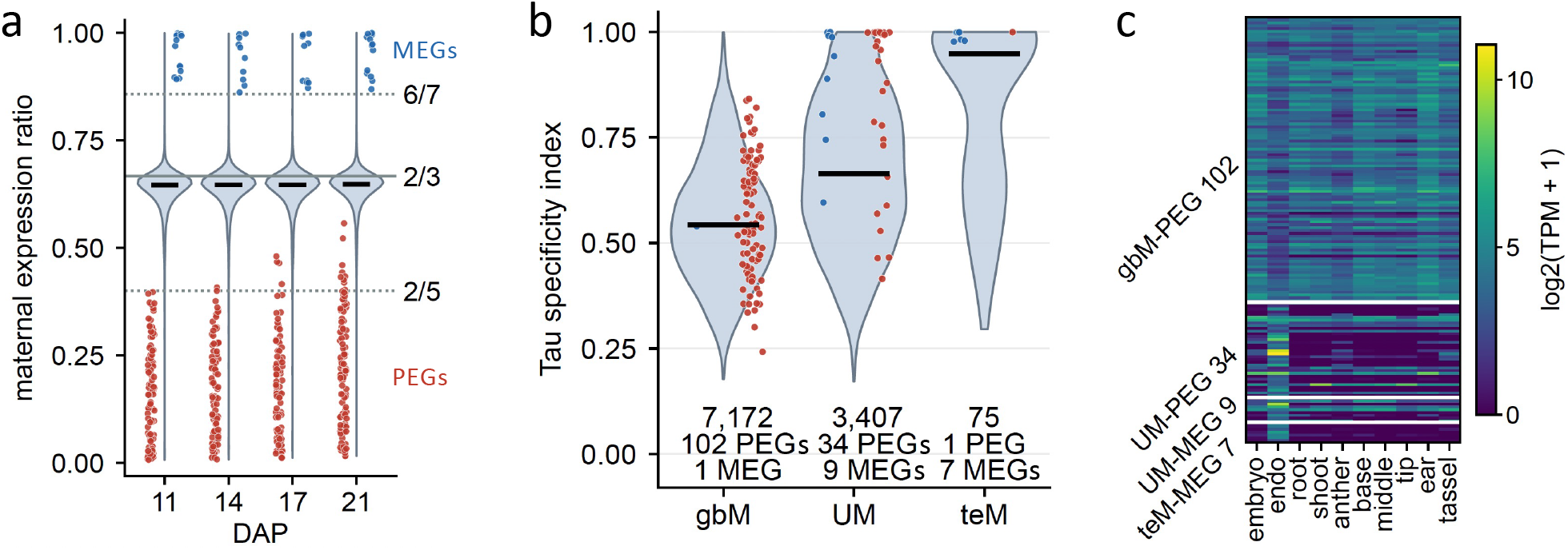
Identification of imprinted genes and their methylation categories. **a.** Distributions of maternal expression ratios from B73/W22 hybrid endosperm from bidirectional crosses at four timepoints in endosperm development, showing all genes whose maternal plus paternal expression was at least 5 RPM in each direction of the cross, for one-to-one B73/W22 genes, with the ratio averaged over the two directions. Dotted lines indicate cutoff values used for defining imprinted genes (blue and red dots) using data from eight hybrids at 14 DAP. Horizontal segments indicate median values. **b.** Methylation categories of imprinted genes correspond to expected expression patterns. The Tau specificity index was calculated from the ten maize tissues shown in panel c. Violins represent the 10,654 genes with defined methylation categories and imprinting calls (the 7,172, 3,407 and 75 under them). **c.** UM-PEGs but not gbM-PEGs are preferentially expressed in endosperm (16 DAP). “base”, “middle”, and “tip” are all leaf segments. Each row corresponds to a single gene, ordered by chromosome and position (excluding the single gbM-MEG and teM-PEG).

### The majority of PEGs have gene body methylation and constitutive expression

137 of the 153 PEGs were categorized according to their gene methylation status using published methods with minor modifications [18]. Of those, 34 were unmethylated (UM-PEGs), 102 were gene body methylated (gbM-PEGs), and one was TE-like methylated (teM-PEGs). The remaining 16 PEGs were uncategorized either due to intermediate methylation values or missing data. Of the 17 MEGs in defined methylation categories, nine were UM-MEGs, one was a gbM-MEG, and seven were teM-MEGs. In comparison, of the 9,143 biparentals in defined methylation categories, 2,787 were UM-biparentals, 6,321 were gbM-biparentals, and 35 were teM-biparentals. These numbers correspond to a 108-fold enrichment for teM genes among MEGs relative to biparental genes (41.2% versus 0.38%, Fisher exact p = 4.7e-13) and a 56-fold enrichment relative to PEGs (41.2% versus 0.73%, Fisher exact p = 4.1e-7), consistent with recent findings [20]. PEGs, in contrast, were 12.7-fold enriched for gbM relative to MEGs (74.5% versus 5.9%, Fisher exact p = 4.3e-8). Given that 16 of the PEGs were in undefined methylation categories, 102 gbM-PEGs of 180 total imprinted genes is an underestimate. Nonetheless, they still made up the largest category of imprinted genes, suggesting a connection between imprinting and gene body methylation.

The methylation categories of imprinted genes corresponded with their expression patterns, with gbM genes being constitutively expressed, UM more endosperm specific, and teM the most endosperm specific (Fig. 1b,c). The Tau tissue specificity index of gbM-PEGs was correspondingly lower than UM-PEGs (median Tau 0.57 versus 0.99, p = 1.3e-10). The median endosperm specificity, measured as endosperm TPM over the highest of nine non-endosperm tissues, was 0.19 for gbM-PEGs and 19.4 for UM-PEGs (p = 9.7e-10), even though TPMs were obtained from a single point in endosperm development 16 days after pollination (DAP) [34]. These results are consistent with prior work in rice and maize showing two categories of PEGs, constitutively expressed ones and endosperm specific ones, where the constitutively expressed ones have reduced maternal gene body methylation, and both groups acquire H3K27me3 on maternal copies in endosperm [30,31,36]. Since H3K27me3 marks dynamically repressed genes, typically UM genes, a prediction of these results is that UM-PEGs but not gbM-PEGs should be repressed by H3K27me3 in non-endosperm tissues. To test this, we examined published H3K27me3 data from B73 leaves and developing ears [37]. We found that UM-PEGs but not gbM-PEGs had H3K27me3 in both tissues (leaf: 59.3% of UM-PEGs were enriched for H3K27me3 versus 1% of gbM-PEGs, p = 5.7e-7; ear: 66.7% versus 1%, p = 5.1e-8; two-sided Mann-Whitney U test on per-gene enrichment; Supplemental Fig. S1).

### Gene body demethylation is not a consequence of maternal repression

The enrichment for gbM genes among PEGs relative to MEGs suggests a connection between maternal gene body demethylation and transcriptional repression. It also raises the question of which causes which. For example, maternal PRC2 repression of gbM-PEGs could inhibit their methylation. To test this, we asked whether gene body demethylation was specific to imprinted genes or included biparentally expressed genes. Comparison of CG methylation in gbM genes in non-endosperm tissues relative to endosperm, however, revealed widespread reduction within gene bodies, including in biparental genes in endosperm (Fig. 2a-d). Here and in reference to per-gene measurements, introns are included. The median CG methylation of gbM-PEGs was 36.5% in embryo and 19.6% in endosperm. The median per-gene pairwise CG methylation change was -15.5% (Supplemental Fig. S2). For brevity we refer to this value as delta and measure it in points, in this case, a delta of -15.5 points (with an IQR of -21.4 to -10.9 points). For gbM-biparentals, the delta was -4.5 points (IQR -9.8 to -0.4). All three non-endosperm tissues agreed (embryo, leaf and tassel), with gbM-PEG deltas of -15.5 to -19.0 points and gbM-biparental deltas of -4.5 to -5.1 points. In contrast, control regions generated by shuffling the gbM-biparental coordinates had similar CG methylation values between endosperm and non-endosperm; for example, their delta between embryo and endosperm was -1.0 points (IQR -3.8 to +1.8). We focused on CG methylation because CHG methylation is less informative for gbM genes, which by definition have CG but not CHG methylation in their coding DNA. CHG methylation in gene bodies also has a slight global reduction in endosperm, as previously observed [15]. CHH methylation in gbM-PEGs and UM-PEGs followed similar patterns in endosperm as in other tissues (Supplemental Fig. S3).

**Figure 2:**
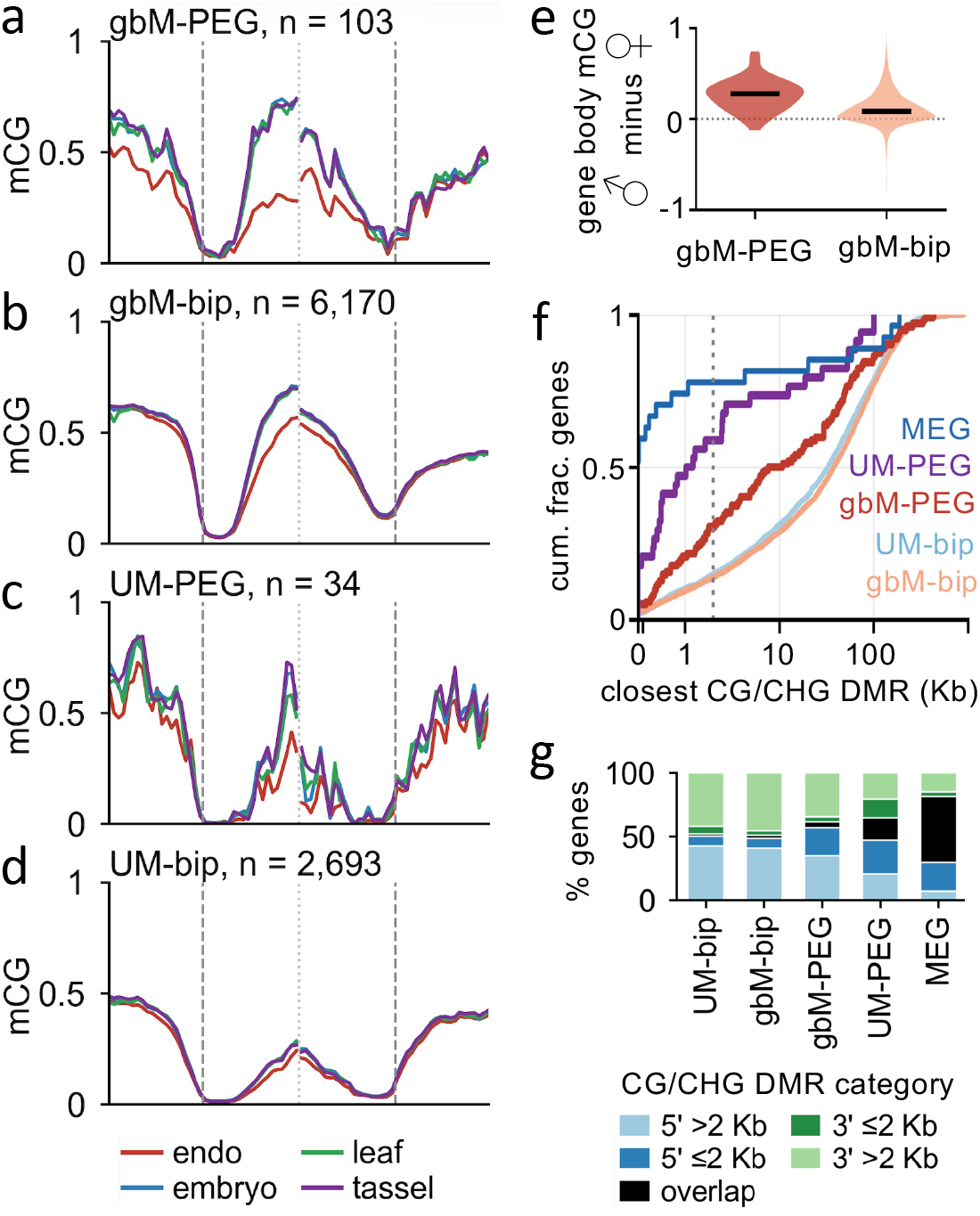
Methylation landscapes of imprinted and biparentally expressed genes. **a-d.** Metagene plots show methylation trends in wild-type W22 15-DAP endosperm and embryo, developing second leaf, and premeiotic tassel. mCG is the fraction of methylated cytosines per total cytosines, calculated on 100-bp intervals, 2 Kb internal and 2 Kb flanking genes on each side. “bip” indicates biparental. Transcription start sites and polyadenylation sites are indicated by vertical dashed lines, mid-gene break by vertical dotted line. “n” indicates largest number of genes with data at any interval **e.** Parentally resolved methylation shows gbM-PEGs and gbM-biparentals have reduced maternal CG methylation in mature W22/B73 hybrid endosperm from bidirectional crosses. mCG was calculated as whole-gene averages. The plot shows the distribution of differences between paternal and maternal syntelogs. Horizontal segments indicate median values. gbM-PEG: n = 76, median +28.0 points; gbM-bip: n = 4,847, median +8.3 points (paternal vs maternal paired two-sided Wilcoxon p = 2.0e-13 and p < 1e-300). **f.** Distances from genes to closest CG/CHG DMR (endosperm demethylated region, defined by comparison between B73 mature endosperm and developing leaf). The dotted vertical line marks 2 Kb. **g.** Location of closest CG/CHG DMR. Stacked bars indicate proportions of genes in each closest DMR location category.

These results raise the question of whether the reduced CG methylation in gbM-biparentals is itself biparental rather than maternal, which could explain both biparental expression and smaller changes in methylation in gbM-biparentals compared to gbM-PEGs. To test this, we examined methylome data from B73/W22 hybrid endosperm [38]. For gbM-PEGs, the median maternal-to-paternal CG methylation delta was 28.0 points (IQR 14.7 to 37.0, n = 76, p = 2.0e-13) (Fig. 2e). While the delta for gbM-biparentals was smaller than that of gbM-PEGs, it was still strongly positive, 8.3 points (IQR 0.4 to 19.2, n = 4,847, p < 1e-300). Measured as a relative difference in methylation (mean maternal value divided by mean paternal value), this corresponds to 24% lower maternal methylation than paternal for gbM-biparentals. These results indicate that maternal demethylation of gbM genes is widespread and not limited to genes with imprinted expression, and thus is not a consequence of maternal repression.

### Cis-regulatory elements vs. gene body demethylation in maternal repression

We focused on gene bodies of gbM-PEGs, but reduced methylation was also evident in their flanks, especially upstream (Fig. 2a). While their core promoters may be constitutively and biparentally unmethylated, enhancers or repressors further upstream could have reduced maternal methylation. If gene body demethylation does not repress gbM-biparental genes, it also may not be what represses gbM-PEGs. Instead, they could be repressed by demethylation of cis-regulatory elements, with gene body demethylation being an inconsequential side effect. Imprinting studies conventionally use demethylated regions (DMRs) selected by comparing WT and DNG mutant endosperm or by comparing WT endosperm and other tissues to identify imprinted genes that are likely explained by demethylation. Since methylation outside of transcribed regions occurs in multiple sequence contexts instead of just CG, canonical imprinting via cis-regulatory elements involves demethylation of multiple sequence contexts. Thus we defined a set of 200-bp DMRs in maize by requiring greater than 50% reduction in both CG and CHG methylation in B73 endosperm relative to leaf using previously described methods [15]. To determine which imprinted gene sets were best explained by these CG/CHG DMRs, we calculated the distance from each set of genes to the nearest ones. MEGs, the best positive control set because of their known imprinting by demethylation of cis-regulatory elements, were closest to DMRs, with 77.8% within two Kb of one (Fig. 2f,g). UM-PEGs also provide a positive control set, but a theoretically weaker one because some have maternal PRC2 recruitment without maternal demethylation [16]. 58.8% of UM-PEGs were within two Kb of a DMR. 30.4% of gbM-PEGs were within two Kb of a DMR (2.6-fold depletion relative to MEGs, p = 1.3e-5). This is still a significant enrichment relative to gbM-biparentals, however, of which only 14.0% were within 2 Kb of a DMR (a 2.2-fold enrichment, p = 2.2e-5). gbM-PEGs were also farther from DMRs than UM-PEGs (median distance 9.2 versus 1.1 Kb, Holm-corrected Mann-Whitney p = 0.003). These analyses are limited by the fact that the DMR set is incomplete, being a representative subset. The results indicate, however, that multi-context demethylation of cis-regulatory elements contributes to maternal repression of an even smaller subset of gbM-PEGs than of UM-PEGs. As discussed below, other evidence suggests that demethylation of cis-regulatory elements without demethylation of gene bodies would be sufficient for activation of gbM-MEGs, but insufficient for repression of gbM-PEGs.

### DNGs mediate maternal demethylation and repression

Demethylation could result as a passive outcome of other chromatin modifications that inhibit maize MET1 methyltransferase activity or by active demethylation by DNGs. To test whether DNGs are required for gene body demethylation, we compared methylation in DNG mutant and wild-type (WT) endosperm. In maize, a pair of DNGs function redundantly in imprinting, MDR1 (DNG101) and DNG102. Since double mutants fail early in embryo development, we analyzed data from single mutants, which have subtle effects on DNA methylation levels because of their redundant function [15,19,38]. In spite of that, each single mutant produced a detectable increase in gene body methylation. In *mdr1* mutant relative to WT developing endosperm (15 DAP), the delta for gbM-PEGs was 4.0 points (IQR 0.8 to 5.9) and for gbM-biparentals was 2.4 points (IQR 0.0 to 5.1) (Fig. 3a). In contrast, shuffled control regions had a delta of 1.2 points (IQR -0.6 to 3.4). In *mdr1* mutant mature endosperm, the delta for gbM-PEGs was 4.2 points (IQR 1.1 to 6.4) and for gbM-biparentals was 2.6 points (IQR -0.2 to 5.7) (Supplemental Fig. S4). In *dng102* mutant mature endosperm, the delta for gbM-PEGs was 4.0 points (IQR 1.0 to 6.6) and for gbM-biparentals was 2.8 points (IQR -0.2 to 6.0) (Supplemental Fig. S4).

**Figure 3:**
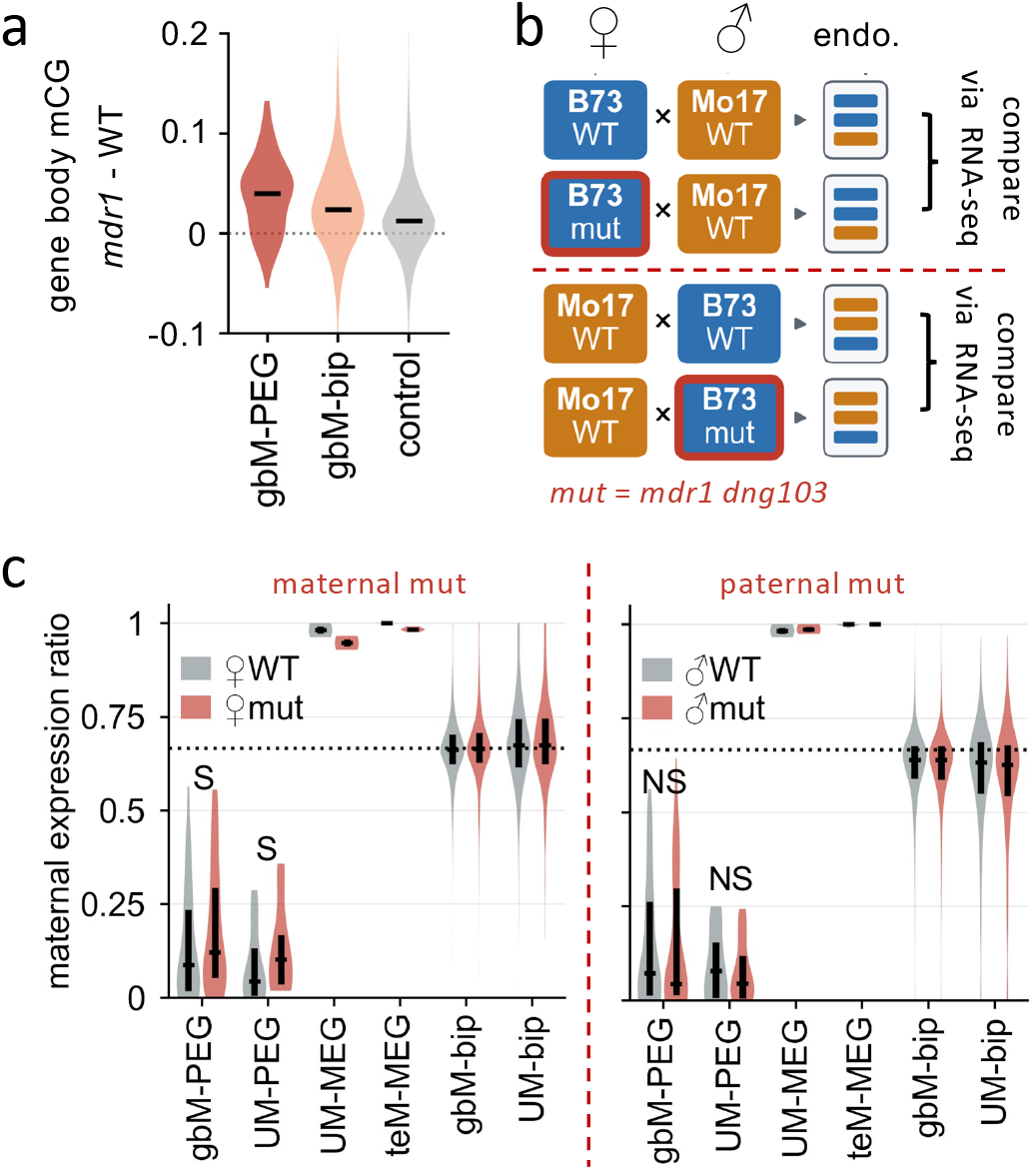
DNGs mediate maternal demethylation and repression. **a.** gbM-PEGs and gbM-biparentals have elevated methylation in *mdr1* single mutant developing endosperm (15 DAP). mCG is the fraction of methylated CGs relative to total CGs calculated as whole-gene averages. The plot shows the distribution of differences between *mdr1* and WT (44 gbM-biparental and 90 control values lie outside the y axis range). Horizontal segments indicate median values. Control regions are shuffled from the gbM-biparentals and have the same distribution of region lengths. **b.** Schematic of experiments for source data in c, from bidirectional B73/Mo17 crosses with B73 parents either WT or *mdr1 dng103* double mutant (“mut”). **c.** Distributions of maternal expression ratios from 14-DAP hybrid endosperm with *mdr1 dng103* double mutant contrasted with WT. The dotted line indicates the expected maternal ratio of 2/3. Horizontal segments indicate medians and vertical segments indicate IQRs. “S” indicates p < 0.001, “NS” p > 0.05, for paired Wilcoxon p-values for differences in PEGs between maternal WT and mutant. 55 gbM-PEGs, 16 UM-PEGs, 2 UM-MEGs, 6 teM-MEGs, 2,834 gbM-biparentals, and 943 UM-biparentals with parentally resolved RNA-seq reads, at least 20 allele-assigned read pairs, summed over the three replicates in both sets of crosses were included in this analysis. The same numbers of genes were included for paternal WT and mutant, except gbM-biparentals decreased to 2,831 and UM-biparentals to 942.

While these results indicate demethylation of gbM-PEGs requires DNGs, they do not answer the question of whether their imprinted expression requires DNGs. Unlike double mutants of *mdr1 dng102*, a double mutant of *mdr1 dng103* produces viable endosperm with a similarly weak methylation phenotype as the *mdr1* single mutant (likely related to the predominant expression of *dng103* in embryo rather than in endosperm) [15,36]. In the prior study of a *mdr1 dng103* double mutant (under the name *zmros1a* for *dng103* and *zmros1b* for *mdr1* in that study), bidirectional crosses produced higher maternal expression ratios of PEGs and lower maternal expression ratios of MEGs in *mdr1 dng103* double mutant. These experiments made use of B73/Mo17 hybrid endosperm from reciprocal crosses with the B73 parent either double mutant for *mdr1* and *dng103* or WT for both genes, and the Mo17 parent always WT (Fig. 3b). To test whether imprinted expression of gbM-PEGs requires DNGs, we analyzed the RNA-seq data from these experiments. In spite of the small number of PEGs with sufficient expression and SNPs for distinguishing parental alleles, gbM-PEGs had significantly higher maternal expression ratios when maternally inherited from a mutant compared to WT (Fig. 3c). Their mutant-to-WT median maternal ratio delta was 2.76 points (IQR -0.60 to 6.13, p = 5.4e-4, paired Wilcoxon). UM-PEGs behaved similarly, with a delta of 3.68 points (IQR 1.38 to 6.45, p = 1.53e-4), as expected if the majority depend on demethylation for maternal repression. Confirming the quality of the source data, the delta for teM-MEGs was in the opposite direction: -1.65 points (IQR -1.84 to -1.32, p = 0.031, and all six genes moved in the same direction). Only two UM-MEGs passed filters, and one moved up and one down. In contrast, gbM-biparentals had maternal expression ratios that changed only negligibly between mutant and WT, though the shift is statistically significant given their large numbers: the per-gene median delta was 0.19 points (IQR -1.66 to 2.26, p = 1.99e-6). Confirming the maternal effect of DNGs, the genotype of the paternal plant had no significant effect on PEGs or MEGs (Fig. 3c). These results indicate that DNG activity causes maternal repression in gbM-PEGs rather than being a consequence of it.

Since we selected gbM-PEGs based on large differences in maternal expression ratios, at least threefold more than expected based on genome dosage, we wondered if the gbM-biparental group was subject to a widespread but more subtle maternal repression by gene body demethylation. To test this, we calculated maternal expression ratios in WT plants for all UM, gbM, and teM biparental genes using data from B73/W22 hybrids at four stages of endosperm development [33]. At all four stages, gbM-biparentals had slightly lower maternal expression ratios than UM-biparentals (medians 0.644 to 0.648 vs 0.650 to 0.651, p = 1.0e-27 to 8.9e-13) (Supplemental Fig. S5). This is consistent with subtle but widespread maternal repression resulting from gene body demethylation, but interpretation is complicated by unmethylated genes also showing lower than the expected maternal expression ratio of 2/3. Too few teM genes were detected (15 to 16) for a meaningful comparison.

### Gene body demethylation is insufficient for H3K27me3 recruitment

Biparental expression of gbM genes could be explained by H3K27me3 having a weak effect on expression or by lack of H3K27me3. To distinguish between these possibilities, we reanalyzed H3K27me3 ChIP-seq data from hybrid endosperm where maternal and paternal H3K27me3 can be resolved by their alleles [32]. Neither the strongly demethylated gbM-biparentals nor the entire gbM-biparental set acquired maternal H3K27me3 in endosperm; gbM-PEGs were distinct from biparentals in this regard (Fig. 4a-e). In particular, their median difference in H3K27me3 enrichment between maternal and paternal syntelogs was 1.34 (IQR 0.25 to 5.96, p = 8.5e-6) (Fig. 4f-j). For strongly demethylated gbM-biparentals, the median difference in H3K27me3 enrichment between maternal and paternal alleles was 0.00 (IQR -0.28 to 0.30, p = 0.76). These were defined as the subset of gbM-biparentals whose CG methylation deltas were in the upper quartile, of at least 16.7 points in the maternal-to-paternal comparison. These were more strongly demethylated than gbM-PEGs, with a median delta of 27.3 points, compared to 23.5 points for the gbM-PEGs used in this analysis. These results indicate that DNA demethylation leads to PRC2 recruitment in PEGs specifically, supporting DNA demethylation as functioning upstream of PRC2. They also indicate that demethylation is not sufficient for H3K27me3 acquisition. As a side note, UM-PEGs had 3’ biased maternal H3K27me3 in endosperm, which we speculate is related to their more frequent 3’ DMRs than either MEGs or gbM-PEGs (Fig. 2g).

**Figure 4:**
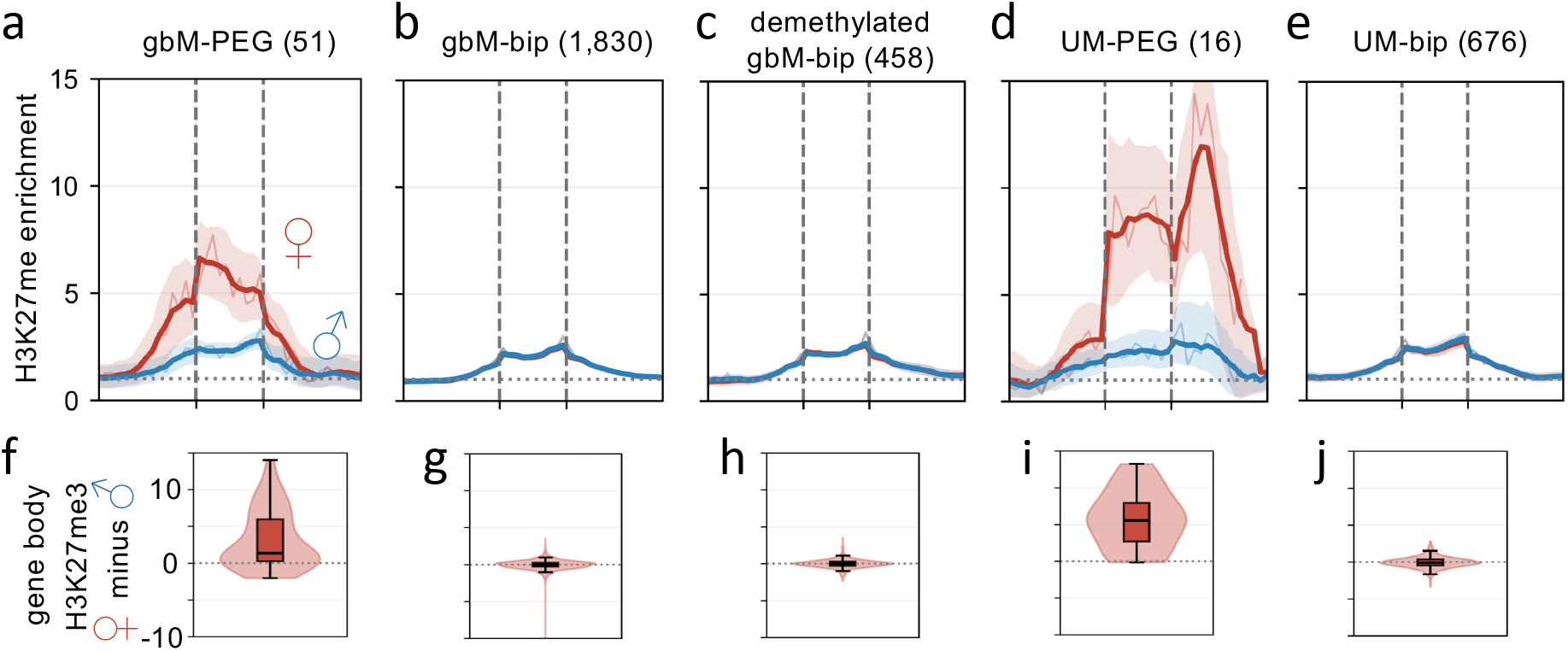
Gene body demethylation is insufficient for H3K27me3 recruitment. **a-e.** Metagene plots of parentally resolved H3K27me3 from ChIP-seq of B73/Mo17 hybrid endosperm from bidirectional crosses. H3K27me3 enrichment is the number of uniquely mapping reads relative to the genome average. Flanks are divided into 1-Kb intervals and extend 15 Kb from transcription start sites and polyadenylation sites, indicated by vertical dashed lines. “demethylated gbM-bip” represent the quarter of gbM-biparentals with the most negative maternal minus paternal mCG, 458 of 1,830, at -16.7 points or lower. Dotted horizontal lines indicate genome average (enrichment value of 1). Internal bins are scaled by gene length: each gene is divided into 10 equal bins. Thick lines indicate a 5-bin centered rolling mean, and thin lines indicate the unsmoothed per-bin values. Shaded areas indicate 95% bootstrap confidence intervals from 1,000 resamples of genes, smoothed over 5 bins. **f-j.** Violins show distribution of maternal minus paternal H3K27me3 enrichments in gene bodies for each set of genes. Values were calculated as whole-gene averages. Whiskers extend to 1.5 × IQR. A single gbM-biparental value is below the axis in panel g.

### Maternal gene body demethylation is the predominant form of imprinting in rice but not in arabidopsis

While it is clear that maternal gene body demethylation is connected to maternal H3K27me3 and repression in rice [30], it is not clear whether this is the most common form of imprinting in rice like it is in maize. Using the imprinted genes that the same study identified in early endosperm development (3-7 DAP) and raw leaf methylome data from a prior study [39], we categorized rice imprinted genes according to their methylation. Of 238 PEGs, 200 had defined methylation categories–41 UM-PEGs, 150 gbM-PEGs, and 9 teM-PEGs (Supplemental Dataset 2). Of 75 MEGs, 60 had defined methylation categories–15 UM-MEGs, 8 gbM-MEGs, and 37 teM-MEGs. These numbers correspond to a 5.6-fold enrichment for gbM genes among PEGs relative to MEGs (two-sided Fisher’s exact p = 5.7e-18) and indicate that gbM-PEGs make up the largest category of imprinted genes in rice. Consistent with our results in maize and an earlier study in rice [40], reanalysis of the endosperm methylome data from the same study that produced lists of imprinted genes [30] showed that maternal CG methylation in gene bodies was lower than paternal in rice gbM-biparentals (Supplemental Fig. S6). Measured as a relative difference in methylation (mean maternal value divided by mean paternal value), rice gbM-biparentals had 26% lower maternal methylation than paternal.

While gene body methylation has not been implicated in imprinting in arabidopsis, PRC2 is strongly implicated in imprinting of arabidopsis PEGs [16]. We asked whether maternal gene body demethylation also occurs in arabidopsis and whether it is common among imprinted genes. We identified 36 PEGs and 70 MEGs by comparison of three published lists of imprinted genes, including only those which were called by at least two of the three [41–43] (Supplemental Dataset 3). 35 of the PEGs and 69 of the MEGs were previously categorized by their methylation status [22]. These were made up of 23 UM-PEGs, 11 gbM-PEGs, and 1 teM-PEG; and 50 UM-MEGs, 6 gbM-MEGs, and 13 teM-MEGs. These numbers correspond to a 3.6-fold enrichment for gbM genes among PEGs relative to MEGs (two-sided Fisher p = 0.0049), but make up a small category of imprinted genes relative to UM-PEGs or UM-MEGs. Reanalysis of hybrid DNA methylation data in arabidopsis [44] confirmed that gbM-PEGs are partially maternally demethylated. The median paternal relative to maternal difference in CG methylation for gbM-PEGs was a delta of 4.6 points (IQR 2.5 to 8.7) (Supplemental Fig. S7e). The change in methylation between paternal and maternal for gbM-biparentals was even more subtle, with a delta of 2.4 points (IQR 0.5 to 4.9). Measured as a relative difference in methylation (mean maternal value divided by mean paternal value), arabidopsis gbM-biparentals had 12% lower maternal methylation than paternal.

To test whether DEMETER (DME), the primary DNG in imprinting, was required for maternal demethylation, we reanalyzed hybrid endosperm methylomes from a cross between a heterozygous *dme* mutant and WT plant [45]. We found that maternal CG methylation was partially restored to paternal levels in the presumed *dme* maternal mutant endosperm compared to endosperm from a WT x WT cross (Supplemental Fig. S7f). The paternal relative to maternal difference in CG methylation for gbM-PEGs yielded a delta of 2.8 points (IQR 0.5 to 9.0) for *dme*, but 9.9 points (IQR 5.1 to 30.6) for WT endosperm (paired n = 8, p = 0.055, not significant). The median paternal relative to maternal difference in CG methylation for gbM-biparentals yielded a delta of 0.0 points (IQR -6.3 to +6.5) for *dme*, but 0.2 points (IQR -3.2 to +9.6) for WT (p = 2.8e-34). These results indicate that maternal gene body demethylation is widespread, but weaker and has a smaller impact on imprinted gene expression in arabidopsis endosperm than in rice or maize.

## DISCUSSION

These findings connect two mysteries of epigenetics: genomic imprinting and gene body methylation. Given its widespread occurrence independently of imprinted expression and maternal H3K27me3, gene body demethylation is not merely a consequence of imprinting. Likewise, since it requires DNGs, it is active, not passive demethylation. What the impact of gbM is on most genes remains debatable because of the lack of clearly interpretable experimental data, the lack of identified methyl-binding proteins to read gene body methylation and translate it to other nuclear functions, and its apparently dispensable nature. The existence of gene body methylation in any tissue strongly corresponds to both gene body methylation and moderate levels of gene expression in other tissues as well. For example, gbM epialleles defined by methylation in early developing leaves in maize are weakly but positively correlated with gene expression levels in nine of ten tissues examined, the single exception being endosperm [18]. This previously unexplained result may be a result of reduced gbM in endosperm and gbM having positive effects on expression in endosperm. In light of our findings and of previous studies showing that constitutively expressed PEGs have maternal specific H3K27me3 in endosperm [30,31], we conclude that gbM functions in protection of a small number of genes from PRC2 repression in endosperm. Since only the maternal copies are demethylated, the paternal copies stay protected.

This interpretation raises the question of how most genes could be subject to maternal gene body demethylation without PRC2 recruitment. A theoretical explanation is that the repressed genes have nearby or internal sequences that can function effectively as endosperm polycomb response elements (PREs) when methylation is removed, while biparental genes lack them (Fig. 5). How would DNA methylation inhibit PRC2? One possibility is through its inhibition of H2A.Z recruitment and maintenance [46,47]. Since H2A.Z in a ubiquitinated form recruits PRC2, the pathway from DNA demethylation to H3K27me3 in endosperm may require ubiquitinated H2A.Z as an intermediate step [48,49]. While the mechanism behind methylation-mediated PRC2 inhibition may not be clear, multiple lines of evidence indicate it happens [26–29].

**Figure 5:**
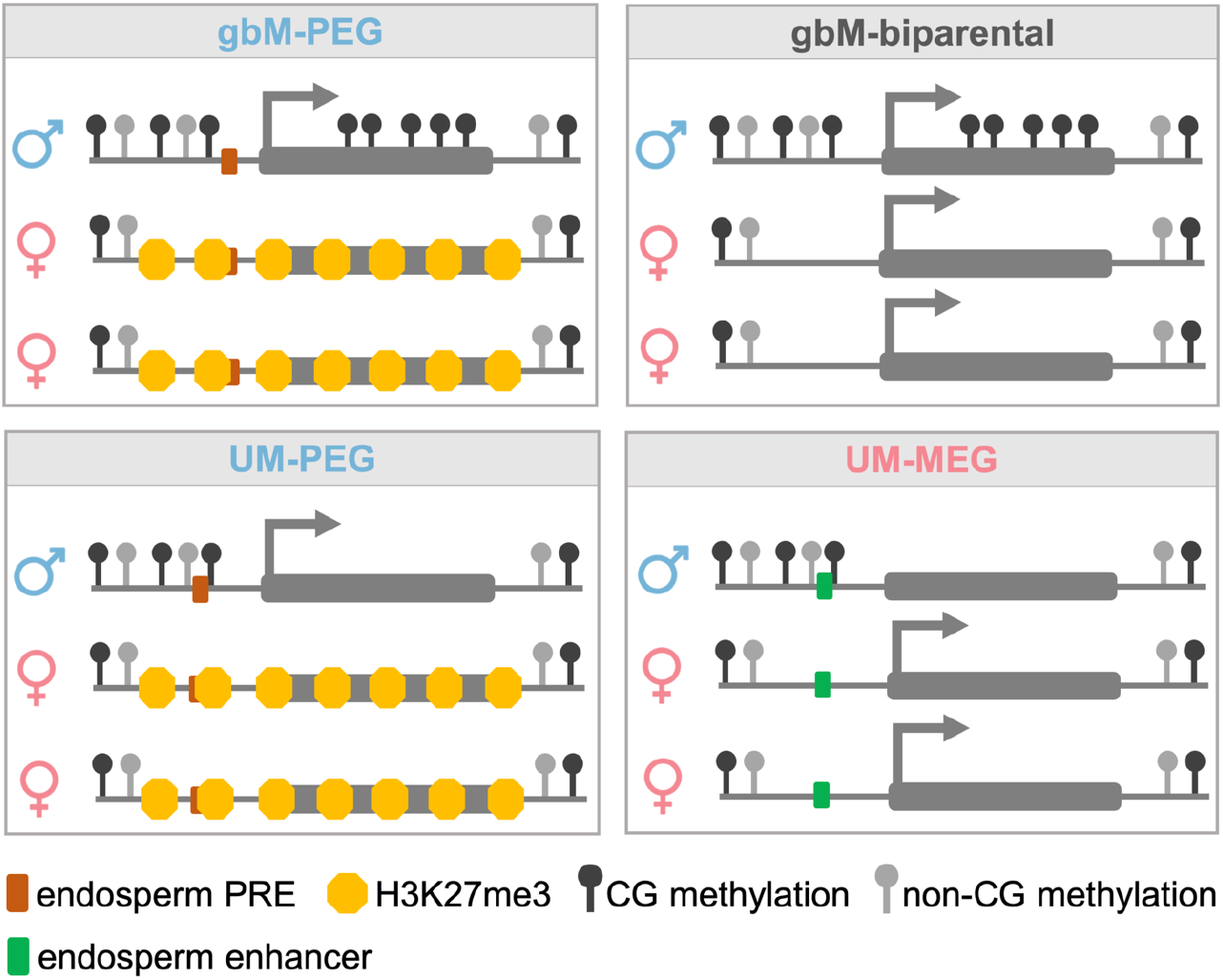
Hypothetical mechanisms of imprinted and biparental expression. Maternal demethylation of gene bodies allows Polycomb Response Elements (PREs) to recruit H3K27me3 and repress genes with CG methylation in their gene bodies (gbM-PEGs) via endosperm trans-acting factors that bind the PREs. In the example, the PRE is located in a constitutively unmethylated region, but it could also be in a demethylated region either flanking the gene or in the gene body itself. In contrast, lack of endosperm PREs would explain why the majority of demethylated gbM genes are biparentally expressed (gbM-biparentals). Genes with constitutively unmethylated gene bodies (UM-PEGs) would be repressed by endosperm PREs much like gbM-PEGs, but by definition would not require demethylation of gene bodies. In some cases they could have demethylated endosperm PREs (as shown in the example). In others they could have constitutively unmethylated PREs but depend on endosperm-specific trans-acting factors for repression rather than on demethylation. MEGs, whether constitutively unmethylated (UM-MEGs) as shown, or TE-like methylated (teM-MEGs), require demethylation of activating cis-regulatory elements.

The fact that demethylation of gene bodies is frequently coupled with demethylation of upstream regions raises the question of whether maternal repression could be explained entirely by demethylation of PREs, which would be more consistent with the conventional understanding of DNGs in imprinting. Two pieces of theoretical evidence support gene body demethylation itself, not just PRE demethylation, being a requirement for imprinting: First, the fraction of gbM imprinted genes is 12.7-fold higher for PEGs (102/137) than MEGs (1/17). This result requires that gbM genes be 12.7-fold more likely to carry maternally demethylated PREs than maternally demethylated transcriptionally activating cis-regulatory elements, which would be difficult to explain. Second, H3K27me3 functions through forming large repressive domains that spread through transcribed regions, not just cis-regulatory elements [25]. If DNA methylation inhibits H3K27me3 recruitment [26–29], demethylation from cis-regulatory elements alone would not allow for H3K27me3 recruitment across the large transcribed regions needed for repression. gbM-MEGs, in contrast to gbM-PEGs, could be explained entirely by demethylation of cis-regulatory elements because MEGs do not require H3K27me3.

A third source of evidence for gene body demethylation as opposed to cis-regulatory element demethylation being required for maternal repression comes from the lower frequency of detectable CG/CHG DMRs in and near gbM-PEGs compared to MEGs, the set that has the strongest evidence for imprinted expression depending on demethylation. Outside of transcribed regions, demethylation of regulatory elements in multiple sequence contexts, not just CG, is what characterizes imprinted genes. The lower frequency of detectable CG/CHG DMRs in and near gbM-PEGs relative to UM-PEGs provides further evidence for gbM-PEGs requiring demethylation in their gene bodies, but with the limitation that UM-PEGs may serve as a weaker comparison than MEGs because many UM-PEGs may be imprinted by mechanisms that do not rely on maternal DNA demethylation [16]. While it would not be surprising for some gbM-PEGs to require demethylation of cis-regulatory elements, we conclude from these results that demethylation of their gene bodies is required for H3K27me3 acquisition and repression of the majority of gbM-PEGs–and further, explains the majority of imprinted genes in maize and close to half in rice.

What targets DNGs to gene bodies? Outside of endosperm, many gene bodies in arabidopsis are already known to be targeted by DNGs in a dynamic balance between methylation and demethylation [50,51]. Multiple lines of evidence indicate that the local chromatin environment and DNA accessibility can control recruitment of DNGs (though not removing the possibility of sequence-specific elements as well) [52–55]. A chromatin state associated with gene bodies may attract DNGs in certain cell types more than others. If, for example, the central cell (the female gamete that gives rise to endosperm) were such a cell and the sperm cell were not, this would result in maternal demethylation in endosperm. Why the effect is strong in rice and maize but weak in arabidopsis could be explained, theoretically, by physiology of their gametophytes or by the amounts of heterochromatin in their genomes.

Genomic imprinting may be a useful method for optimizing gene expression levels in endosperm that relies on the chromatin differences inherited from central cell and sperm [56]. But what advantage would epigenetic control offer compared to genetic control of gene expression? One is its variability, not just from individual to individual, but even from cell to cell. In fact, Kermicle’s original imprinting phenotype was called mottling because of its cell-to-cell variation in pigmentation [5]. Related to this, a possible consequence of widespread gene body demethylation in endosperm independent of imprinting is increased gene expression variability [24,50]. Theoretically, variability in gene expression is beneficial as a source of bet hedging, where randomized subsets of cells are optimized for different conditions [57]. The best studied examples of this in plants are related to seed dormancy and germination time. If gene body methylation indeed reduces noise in endosperm, its demethylation would increase noise for a large number of genes independently of imprinting and increase noise for a small number through imprinting.

## DATA AVAILABILITY

No new sequencing data were generated for this study. All analyses used published data. Processed files from publication or public repositories are listed in Supplemental Table S1. The sources of all raw sequencing reads reanalysed from public repositories are listed in Supplemental Table S2. All code, with conda environment files and a README that maps each figure panel and each number in the text to the script that produced it, is available at https://github.com/chiragh241/gbm-imprinting. Supplemental Table S3 lists the scripts that produce each figure.

## METHODS

### Software sources

All processing ran under WSL2 (Ubuntu 26.04 LTS) in five conda environments (envs/). Read processing used Python 3.10.20, methylpy 1.4.7 [58], Bismark 0.24.2 [59], bowtie2 2.5.5 [60], cutadapt 5.2 [61], Picard 3.4.0 [62], bedtools 2.31.1 [63], samtools 1.23.1 [64], sra-tools 3.4.1 (https://github.com/ncbi/sra-tools), HISAT2 2.2.1 [65], featureCounts 2.0.8 (Subread) [66], minimap2 2.28 [67] and SNPsplit 0.6.0 [68]. BS-Seeker2 2.1.7 [69] ran on Python 2.7.15 with bowtie2 2.5.4, samtools 1.18 and pysam 0.20.0, and CGmapTools 0.1.3 [70] (commit afa6e40) with samtools 1.22.1 and R 4.6.0. Imprinting calls, downstream analysis, statistics and figures used R 4.5.1 and Python 3.13.7 with pandas 2.2.3, NumPy 2.2.6, SciPy 1.16.2 and Matplotlib 3.10.6.

### Identification of maize imprinted genes

Imprinted genes were called from allele-resolved endosperm RNA-seq of B73/NAM-founder and B73/W22 reciprocal hybrids as summarized for each gene and hybrid in the table B73_NAM_syntelog_maternalpref.txt. This table contains 237,338 of the 444,878 rows of the published table mat_pref_NAM.txt [33] with two additional features added for each row: “feature_2” is the B73 v5 syntelog(s), and “cont_genome” is the code name for the hybrid as defined in the prior study. (The two-letter names for each hybrid are as follows: BB = B97, BK = Ki11, BO = Oh43, BW = W22, CB = NC358, DB = Ky21, EB = CML333, and FB = M162W. Data from the other B73/Oh43 hybrid used in the study were excluded). Each row reports allele-specific expression in RPM for one syntelog pair in one hybrid in one direction of the cross obtained by mapping reads uniquely to the non-B73 (type A) and B73 (type B) reference genomes. RPM values are reported as A means and B means, since each RPM value was originally obtained with biological replicates. For B73 paternal crosses, the maternal ratio of each syntelog pair is computed as the A mean divided by the sum of the A and B means. For B73 maternal crosses, the maternal ratio is the B mean divided by the sum of the A and B means. Thus each syntelog pair produced two maternal ratio values. The retained 237,338 rows from mat_pref_NAM.txt are those whose A and B means sum to at least two RPM and have a B73 v5 syntelog. This rule, reconstructed after the fact, recovers all 237,338 rows but would admit 848 more, so the table is provided in the repository as used (data/maize/B73_NAM_syntelog_maternalpref.txt.gz). Rows whose gene is linked to several B73 genes were expanded into one row per B73 syntelog before calling (01_imprinting_calls/01_identify_imprinted.R).

Since the null expectation for a biallelically expressed gene is a 2:1 maternal:paternal ratio (maternal ratio of 2/3, about 0.667), genes were called imprinted when allelic expression deviated at least 3-fold from this expectation. This yielded 27 maternally expressed genes (MEGs) with maternal ratios of at least 6/7 (about 0.857), 153 paternally expressed genes (PEGs) with maternal ratios of at most 2/5 (0.40), and 11,331 biparental genes with maternal ratios between these cutoffs. In making these imprinting calls, five filters were applied to reduce false positives from gene expression variation. First, the total expression (sum of the A and B means) for each record had to be at least five RPM to exclude unreliable ratios from low coverage. Second, the sum of the A and B means had to be at least one-third of the average total for that gene across hybrids in the same direction of the cross. This removed measurements in which a gene was weakly expressed in a particular hybrid background, where low counts produce noisy ratios that can make an imprinted gene appear biparentally expressed. Third, for each syntelog pair in a hybrid cross, both directions of the cross had to yield the same imprinting call. Fourth, each gene had to have consistent calls from at least two hybrids. Fifth, each gene had to have consistent imprinting calls from all hybrids that produced imprinting calls. The 13,209 genes with at least one hybrid passing the first three filters are referred to as evaluable for imprinting; besides the 27 MEGs, 153 PEGs and 11,331 biparental genes, 1,587 had a call from only one hybrid (insufficient data) and 111 had conflicting calls between hybrids (unknown). Per-measurement calls (one row per gene x hybrid x alignment type), a per-gene collapsed table, and the full genome-wide call sheet (including biparental, insufficient-data, and unknown genes) were written to imprinted_3fold_genes.csv, imprinted_3fold_genes_ collapsed.csv, and b73_imprinting_calls_all.csv, respectively. Imprinting calls were generated with an R script (01_imprinting_calls/01_identify_imprinted.R) using R 4.5.1 and the tidyverse packages readr 2.2.0, dplyr 1.2.1, tidyr 1.3.2 and stringr 1.6.0. All thresholds (MEG cutoff 6/7, PEG cutoff 2/5, minimum rpm_sum 5, expression-outlier fold 3, minimum 2 hybrids) are defined as named constants at the top of the script. Calls for every evaluable gene are given in Supplemental Dataset 1 (06_per_gene_methylation/01_supplemental_dataset1.py).

Maternal expression ratios across endosperm development (Fig. 1a, Supplemental Fig. S5) were taken from the reciprocal B73 x W22 time course of Higgins et al. (B73_W22_maternalpreference_with_avg_rpm.csv) [33], which gives the RPM of each allele at 11, 14, 17 and 21 DAP in both directions of the cross. In each direction the maternal ratio is the maternal allele’s RPM divided by the sum of both alleles: the B73 allele in B73 x W22 and the W22 allele in W22 x B73; the recomputed ratios reproduce the file’s own ratio columns. The two directions list genes under B73 and W22 identifiers respectively, so they were paired through b73v5_to_w22.tsv, keeping one-to-one pairs only (12,099 genes). At each stage a gene was used when both alleles summed to at least 5 RPM in each direction, and its maternal ratio is the mean of the two directions’ ratios, which cancels differences between the B73 and W22 alleles (lib/timecourse.py). Fig. 1a plots every gene passing this filter, 7,352 to 8,680 per stage, with MEG and PEG calls marked (12_figures/01_fig1.py). Supplemental Fig. S5 plots the same averaged ratios for biparental genes by methylation category and compares gbM with UM genes at each stage by two-sided Mann-Whitney U test (12_ figures/13_figS4.py).

### Gene expression specificity calculations

Expression in ten B73 tissues was taken from the per-gene TPM values of Zeng et al. [18], computed there from the NAM RNA-seq of Hufford et al. [34]: embryo and endosperm at 16 DAP, root and shoot (coleoptile) six days after planting, base, middle and tip of the tenth leaf at V11, meiotic tassel and immature ear at V18, and anther at R1. The values were read from b73.all.6_38.txt, whose ten source-tissue columns are those of B73.all.csv of Zeng et al. [18]. Two genes appear twice in that table under different pan-gene identifiers with identical values (Zm00001eb007510 and Zm00001eb390120) and were kept once. Tissue specificity was quantified per gene with tau over the ten tissues, tau = sum over i of (1 - x_i / x_max) / (n - 1) with n = 10, which runs from 0 for a gene expressed equally in every tissue to 1 for a gene expressed in one. Negative values would be set to zero, and genes with no expression in any tissue have no tau and were dropped (12_figures/01_fig1.py). Fig. 1b shows the 10,654 evaluable genes that carry a gbM, UM or teM category. Categories were compared by Kruskal-Wallis test, and gbM-PEGs with UM-PEGs by two-sided Mann-Whitney U test. Endosperm specificity was computed per gene as endosperm TPM divided by the highest TPM among the other nine tissues. A gene expressed in endosperm and nowhere else has an infinite ratio; it was kept and ranked above every finite value, which leaves the median and rank-based tests well defined (03_expression_specificity/01_endosperm_ specificity.py). gbM-PEGs and UM-PEGs were compared by two-sided Mann-Whitney U test. Fig. 1c shows log2(TPM + 1) for each imprinted gene in each tissue, genes grouped by category and ordered by chromosome and position within each group (12_figures/01_fig1.py).

### Assignment of maize genic DNA methylation categories

Each gene was assigned a DNA methylation category independently of its imprinting status, from the per-gene CDS methylation values of Zeng et al. [18] (B73.all.csv). mCG and mCHG are the methylated fraction of all CDS cytosines of the gene in each context, cCG and cCHG the numbers of covered CG and CHG sites. Categories were assigned with the rules of Zeng et al. (their GeneMethylationType.sh), in its order and with its inclusive bounds (02_gene_sets/01_categorize_methylation.py): a gene with at least 10 covered CG and 10 covered CHG sites was UM if mCG and mCHG were both at most 0.05, otherwise gbM if mCG was at least 0.2 and mCHG at most 0.05, otherwise teM if mCG and mCHG were both at least 0.4. Genes with fewer covered sites, meeting none of the three definitions, or absent from the table were labelled undefined; 31,654 of the table’s 39,756 genes received a category (15,716 UM, 9,090 gbM and 6,848 teM). The rules differ from those of Zeng et al. only in the coverage minimum, which had been set at 40 CG and 40 CHG sites; 40 would leave four of the seven teM-MEGs undefined. Applied at 40 sites, the same rules reproduce the published categories of Zeng et al. for every gene. Overlapping the four imprinting categories (MEG, PEG, biparental, and undefined) with the four methylation categories gave 16 mutually exclusive sets over the 13,209 evaluable genes (02_gene_sets/ 03_build_sets.py). In B73 v5 these comprise 102 gbM-PEGs, 34 UM-PEGs, 1 teM-PEG and 16 undefined PEGs; 1 gbM-MEG, 9 UM-MEGs, 7 teM-MEGs and 10 undefined MEGs; and 6,321 gbM, 2,787 UM, 35 teM and 2,188 undefined biparental genes. The 1,698 genes without an imprinting call were 748 gbM, 577 UM, 32 teM and 341 undefined. Enrichment of a methylation category in one imprinting category relative to another was tested by two-sided Fisher’s exact test on genes with a methylation category (02_gene_sets/04_enrichment_stats.py). Gene coordinates were taken from the MaizeGDB B73 v5 annotation (Zm-B73-REFERENCE-NAM-5.0_Zm00001eb.1.gff3). For analyses of W22 data, each B73 gene was mapped to its W22 syntelog(s) through b73v5_to_w22.tsv, which pairs every B73 v5 gene with all W22 v2 genes (“Zm00004b” prefix) in its row of the MaizeGDB 2020 pan-gene table (Supplemental Table S1) (02_gene_sets/02_b73v5_to_w22.py), with W22 coordinates from the W22 v2 annotation [35]. A B73 gene may have several W22 syntelogs or none: 514 genes have none and drop out, and a gene with more than one contributes one interval per syntelog, giving 12,842 W22 intervals. In W22 the sets are 103 gbM-PEGs, 34 UM-PEGs, 1 teM-PEG and 15 undefined PEGs; 1 gbM-MEG, 7 UM-MEGs, 4 teM-MEGs and 10 undefined MEGs; and 6,181 gbM, 2,696 UM, 25 teM and 2,148 undefined biparental genes (721, 556, 20 and 320 without a call). The one extra gbM-PEG is Zm00001eb133500, which has two W22 syntelogs. Because the categories are B73-derived, they were not re-derived in W22.

### Methylation calling, metagene methylation, and per-gene methylation

EM-seq reads of W22 inbred tissues were obtained from BioProject PRJNA759188: wild-type, *mdr1* and *dng102* endosperm, wild-type 15-DAP embryo, developing second leaf and premeiotic tassel, three biological replicates each (24 runs, Supplemental Table S2). Runs were fetched from the SRA open-data mirror, checked with vdb-validate and converted to FASTQ with fasterq-dump (sra-tools 3.4.1) (04_emseq_methylation/01_download_reads.sh). Forward and reverse bisulfite-converted references of the W22 v2 genome were built with methylpy build-reference --aligner bowtie2 (02_bisulfite_reference.sh). Each replicate was processed separately with the methylpy 1.4.7 paired-end pipeline (methylpy paired-end-pipeline --num-procs 8 --remove-clonal True --sort-mem 2G), which trims adapters with cutadapt 5.2 (methylpy defaults: read 1 AGATCGGAAGAGCACACGTCTGAAC, read 2 AGATCGGAAGAGCGTCGTGTAGGGA, minimum read length 30), aligns with bowtie2 2.5.5, removes clonal reads with Picard 3.4.0 MarkDuplicates, keeps alignments with MAPQ at least 30 and writes per-cytosine methylated and total read counts (allc tables) (03_map.sh). methylpy was patched to run with these versions: it no longer passes cutadapt the format option removed in cutadapt 2.0, and it adds a default read group before MarkDuplicates, which Picard 3 requires (envs/patches/methylpy.patch). The three replicates of each genotype and tissue were then summed per cytosine with methylpy merge-allc (04_merge_allc.sh).

Methylation of a region was quantified throughout as the weighted methylation ratio: methylated reads divided by total reads, summed over all cytosines of one context in the region. Contexts were read from the trinucleotide in the allc table: CG if the second base is G, otherwise CHG if the third base is G, otherwise CHH. Cytosines were assigned to W22 gene intervals with bedtools 2.31.1 intersect (06_region_hits.sh). A gene or region was used only when it had at least 5 covered CG sites and at least 10 reads in every sample being compared, applied identically to every gene set and to the control regions. A control set of 6,181 regions was generated by shuffling the W22 gbM-biparental intervals over chromosomes 1 to 10 with bedtools shuffle -noOverlapping -seed 42, otherwise unconstrained, so the control matches that set in number and length but not in sequence composition (05_shuffled_control.sh). It was processed identically to the gene sets and is labelled shuffled control in the figures.

Metagene profiles (Fig. 2a-d) used fixed 100 bp bins: 20 bins upstream of the transcription start site, 20 bins running inward from the TSS, 20 bins running inward from the annotated 3′ end of the gene, and 20 bins downstream, ordered 5′ to 3′ with respect to the gene (07_metagene_bins.sh). An inward bin was used only if it lay wholly within the gene and did not cross the gene midpoint, so no base pair is counted twice and short genes contribute only to the outer inward bins; genes shorter than 200 bp were skipped. The gene interior between the two inward blocks is not represented and is drawn as a break. For each tissue, gene set and bin, methylated and total CG reads were summed over all genes in the set and their ratio taken (08_ metagene_per_gene.sh, 12_figures/02_fig2_metagene_table.py). Each panel shows one gene set (gbM-PEG, gbM-biparental, UM-PEG, UM-biparental) in four tissues (15-DAP endosperm, 15-DAP embryo, developing second leaf, premeiotic tassel); the number of genes reaching each bin is in the source data and the panel title gives the largest (12_figures/03_fig2.py). CHG and CHH profiles (Supplemental Fig. S3) were built on the same bins, genes and estimator (04_emseq_methylation/09_metagene_by_context.py, 12_figures/11_figS2.py).

Changes between tissues were computed per gene as the endosperm value minus the value in the other tissue, for genes passing the coverage filter in both, and summarised as the median of the per-gene differences in percentage points with its interquartile range (IQR); the shuffled control regions were treated the same way (06_per_ gene_methylation/05_tissue_deltas.py). DNG mutants were compared with wild-type only at matched stages: *mdr1* against wild-type in 15-DAP endosperm, and *mdr1* and *dng102* each against wild-type in mature endosperm. For each gene the difference was mutant minus wild-type, summarised the same way (06_per_gene_methylation/02_ mutant_deltas.py). Paired differences were tested by two-sided Wilcoxon signed-rank test, and two gene sets were compared on their per-gene differences by two-sided Mann-Whitney U test with Cliff’s delta. Fig. 3a shows the same 15-DAP per-gene differences for gbM-PEGs and gbM-biparentals, and per region for the shuffled control (06_ per_gene_methylation/03_control_region_deltas.sh, 04_control_deltas_summary.py), plotted as *mdr1* minus wild-type (12_ figures/05_fig3.py). Supplemental Fig. S4 shows the mature-endosperm comparisons (12_figures/12_figS3.py). Per-gene values for every genotype and tissue are given in Supplemental Dataset 1.

### Allele-resolved maize DNA methylation

Reciprocal F1 mature endosperm of B73 x W22 and W22 x B73, three biological replicates each, was taken from PRJNA759188 (Supplemental Table S2). Where a replicate had been sequenced twice, the two runs were concatenated into one library (05_allele_methylation/02_merge_libraries.sh). A bisulfite index of the W22 v2 genome was built with bs_ seeker2-build.py --aligner=bowtie2 (BS-Seeker2 2.1.7 with bowtie2 2.5.4) (03_bsseeker2_index.sh). The reference was not N masked at B73/W22 variants, because masking a C/T variant removes positions that bisulfite reads have to be read through; alleles were resolved from variant calls instead (below). Reads were aligned with bs_seeker2-align.py -- aligner=bowtie2, in pieces of 4 million read pairs whose alignments were concatenated in order; reads are aligned independently, so the split does not change the result (04_map.sh). Alignments were name-collated, mate-fixed, coordinate-sorted and deduplicated with samtools 1.18 (collate, fixmate -m, sort, markdup -r) (05_sort_dedup.sh). For variant calling, the six deduplicated libraries were merged at full depth (06_pool_endosperm.sh) and per-cytosine ATCGmap files of the merged alignments were written per sequence with bs_seeker2-call_methylation.py --sorted (07_ call_methylation.sh). With pysam releases later than 0.7.7 this step silently discards every read, because BS-Seeker2 sets no proper-pair flag and writes no base qualities; we kept unpaired reads and removed the base-quality minimum, restoring the tool’s original behaviour (envs/patches/bs_seeker2-call_ methylation.patch).

B73/W22 single-nucleotide variants were called from the pooled endosperm ATCGmaps with cgmaptools snv (CGmapTools 0.1.3), keeping sites with read depth 15 to 70, the upper bound excluding collapsed repeats; C/T and G/A variants were kept (08_snv.sh). Allele-specific methylation was called for each library with cgmaptools asm -m ass -d 2, which requires at least two reads per allele at a site, separately for CG, CHG and CHH (10_asm.sh). To bound run time it was called at a subset of the variants: the 472,785 lying within the W22 intervals of the 16 gene sets or within two Kb of them, and, as a non-genic baseline, every 93rd of the remaining variants in file order (50,381) (09_asm_snv_subset.py). Maternal and paternal alleles were assigned from the cross direction rather than the genotype. With W22 as the reference, allele 1 is W22 and allele 2 is B73 (checked row by row), so the maternal allele is allele 2 in B73 x W22 and allele 1 in W22 x B73. A cytosine linked to several variants was counted once, keeping its best-covered row. For each W22 gene interval, methylated and unmethylated CG reads were summed per allele over the six endosperm libraries, and a gene was used when it had at least 3 linked CG sites and at least 10 reads on each allele (11_gene_summary.py). Fig. 2e plots per-gene paternal minus maternal CG methylation for gbM-PEGs and gbM-biparentals, tested by two-sided Wilcoxon signed-rank test on the paired maternal and paternal values (12_figures/ 03_fig2.py). Strongly demethylated gbM-biparental genes (Fig. 4) were the quarter of gbM-biparentals with the largest maternal CG loss (maternal minus paternal at or below -16.7 points; 458 of 1,830). For each gene and cross direction, maternal minus paternal CG methylation was computed from the reads of that direction’s libraries when both alleles had at least 10 reads, then averaged over the directions available (10_chip_endosperm_allele/03_allele_ panel.py).

### Closest DMRs

Endosperm CG/CHG demethylated regions (DMRs) were taken as a set of 27,402 B73 v5 regions of 200 bp in which mature endosperm CG and CHG methylation is at least 50% lower than in early developing leaf (B73_endo-leaf_hyperDMRs.bed) using the same method as previously [15] and B73 endosperm EM-seq source reads from the same BioProject (PRJNA759188) and leaf EM-seq reads from the NAM assembly project [34]. Genes on chromosomes 1 to 10, in B73 v5 coordinates, were written to a BED file with their imprinting and methylation categories (07_dmr_distances/01_prep_beds.py) and analysed as MEGs, gbM-PEGs, UM-PEGs, gbM-biparentals and UM-biparentals. For each gene, the nearest DMR was found with bedtools closest - D a -t first (bedtools 2.31.1) (02_closest.sh). Each gene was then placed in one of five zones by the position of that DMR: overlapping the gene, within 2 kb upstream, within 2 kb downstream, more than 2 kb upstream, or more than 2 kb downstream (03_distance_stats.py). Fig. 2f shows the cumulative distribution of absolute distance and Fig. 2g the fraction of genes in each zone (12_figures/03_fig2.py). Each PEG category was compared with the biparental genes of the same methylation category, because gbM and UM genes differ in their distance to DMRs regardless of imprinting. Absolute distances were compared by two-sided Mann-Whitney U test and the fraction of genes overlapping a DMR by two-sided Fisher’s exact test, for gbM-PEG against gbM-biparental, UM-PEG against UM-biparental and gbM-PEG against UM-PEG; these six tests were Holm-corrected together (03_distance_stats.py). The fraction of genes whose nearest DMR lies within 2 Kb (absolute distance at most 2,000 bp, including overlap) was compared between categories by two-sided Fisher’s exact test, uncorrected (03_distance_stats.py).

### Maize RNA-seq read processing, mapping and allele-specific counts

Endosperm RNA-seq of reciprocal B73 x Mo17 crosses, in which the B73 parent was either wild-type or the *zmros1a zmros1b* (*dng103 mdr1*) double mutant [36], was reanalysed from reads (PRJNA739488, 12 runs, three biological replicates of each of four crosses; Supplemental Table S2). Mo17 was wild-type in every cross: the mutation is maternal in *zmros1ab* x Mo17 (compared with B73 x Mo17) and paternal in Mo17 x *zmros1ab* (compared with Mo17 x B73) (data/rnaseq/samplesheet.tsv). B73/Mo17 single-nucleotide variants were called independently of any published variant set: the Mo17 CAU-2.0 genome [71], split into 1 Mb pieces to bound memory, was aligned to the B73 v5 genome with minimap2 2.28 (-cx asm5 - -cs), and variants were called with paftools.js call -L 5000, giving 4,636,099 single-nucleotide variants (08_rnaseq_ allele/02_build_snps.sh). The B73 v5 genome was N-masked at these positions with bedtools maskfasta and indexed with hisat2-build (03_masked_genome_index.sh). Reads were not trimmed. They were aligned with HISAT2 2.2.1 (--no-mixed --no-discordant --no-unal), assigned to the B73 or Mo17 allele with SNPsplit 0.6.0 (--paired), and counted per gene over the exons of the B73 v5 annotation with featureCounts 2.0.8 (-p --countReadPairs -s 0 -F SAF) (04_align_ split_count.sh).

For each gene and cross, maternal and paternal fragment counts were summed over the three replicates, and the maternal ratio (maternal fragments divided by maternal plus paternal fragments) was computed when the sum was at least 20 (05_maternal_fraction.py). For each gene set, the maternal ratio with a mutant mother was compared with the ratio with a wild-type mother over the genes scored in both crosses, as the per-gene difference (mutant minus wild-type) in percentage points, by two-sided Wilcoxon signed-rank test; the two paternal-mutant crosses were compared the same way as the control (06_delta_by_class.py, 12_figures/05_fig3.py). The six gene sets (gbM-PEG, UM-PEG, UM-MEG, teM-MEG, gbM-biparental, UM-biparental) are separate questions and their p values were not corrected across sets.

### ChIP-seq enrichment calculations from B73 leaf and ear

Mapped reads from H3K27me3 ChIP-seq and input of B73 seedling leaf and developing ear were obtained in B73 AGPv4 coordinates (GSE120304; two ChIP and two input libraries for leaf, two ChIP and one input for ear; Supplemental Table S1) (09_chip_leaf_ear/01_download.sh) [37]. B73 v5 genes were placed in v4 coordinates through the MaizeGDB v5-to-v4 gene cross-reference, keeping only genes with exactly one v4 counterpart, with coordinates from the v4 annotation (02_annotation.sh, 03_gene_intervals.py). Replicates were pooled within each tissue. For the whole-gene measures, ChIP and input reads over each gene body were counted with bedtools 2.31.1 intersect (04_count.sh) and converted to RPKM with a pseudocount of one read, and enrichment was log2(ChIP RPKM / input RPKM). Genes with fewer than 10 input reads over the gene body were excluded, since the value is a ratio to input. A gene was called enriched when its ChIP/input ratio was above 2 (05_gene_enrichment.py, 12_ figures/10_figS1.py). The log2 enrichment of UM-PEGs and gbM-PEGs was compared by two-sided Mann-Whitney U test (p = 5.7e-7 in leaf and 5.1e-8 in ear). For the metagene profiles, each gene was divided into 20 upstream bins of 250 bp, 60 body bins scaled to its length and 20 downstream bins of 250 bp, ordered 5′ to 3′ (06_metagene_bins.py), and ChIP and input reads were counted per bin (07_metagene_count.sh). Because the input libraries are shallow (about 0.4x), a 250 bp bin holds well under one input read, so per-gene enrichment in each bin was ChIP reads per bp in that bin divided by the gene’s input reads per bp over its whole window, scaled by the ratio of input to ChIP library sizes (08_metagene_profile.py). Supplemental Fig. S1 plots the mean of this value over genes with at least 10 input reads on a linear scale, with input level at 1, and shows per gene log2 values as heatmaps (12_figures/10_figS1.py).

### ChIP-seq read processing, mapping, enrichment calculations for allele-resolved hybrid endosperm

H3K27me3 ChIP-seq of 12-DAP endosperm from reciprocal B73 x Mo17 crosses [32] (SRP011991; SRR748861 with a B73 mother, SRR748862 with a Mo17 mother) was reanalysed from reads. No input, mock or IgG library exists for these samples. Single-end reads were downloaded as FASTQ from ENA (10_chip_endosperm_allele/01_download_reads.sh), aligned to the N-masked B73 v5 genome described above with HISAT2 2.2.1 (--no-spliced-alignment --no-unal), filtered to MAPQ at least 30 with samtools view - q 30 and coordinate-sorted (02_align_split_count.sh). A 36 bp read can be assigned to an allele only if it covers a variant, so alignments were first restricted to windows running from 35 bp before each of the 4,636,099 B73/Mo17 variants to the variant (merged with bedtools merge), then assigned to alleles with SNPsplit 0.6.0, and counted per gene with featureCounts 2.0.8 (-s 0 -F SAF). Genes were used when they had at least 20 allele-assigned reads in each of the two libraries: 51 gbM-PEGs, 16 UM-PEGs, 1,830 gbM-biparentals and 676 UM-biparentals (03_allele_ panel.py). Strongly demethylated gbM-biparentals are the 458 of the 1,830 in the lowest quartile of maternal minus paternal CG methylation (at most −16.7 percentage points; see allele-resolved methylation above). Metagene bins were built in B73 v5 coordinates with 20 upstream and 20 downstream bins of 1 kb and 10 body bins, each a tenth of the gene, ordered 5′ to 3′ (04_gene_meta.py, 05_compact_bins.py). Each allele-assigned read was counted once, in the bin holding its leftmost aligned base, the position on which the variant windows are defined; allele-informative base pairs per bin were counted as the overlap of the bin with the variant windows (06_compact_ counts.sh). For each gene set and bin, reads were summed over genes and divided by the summed informative base pairs, and each allele was divided by its own genome-wide density of assigned reads per informative base pair, so that 1 is that allele’s genome average and the 2:1 maternal dosage of endosperm divides out; the two reciprocal crosses were then averaged (07_compact_profile.py). Bins with fewer than 30 pooled reads were left blank, and shaded bands are 95% bootstrap intervals over genes, 1,000 resamples. Profiles were computed to 20 kb from the gene and are shown to 15 kb, the first distance at which the maternal and paternal curves of both PEG sets had met on both sides. Values are drawn as a centred five-bin rolling mean (5 kb in the flanks), run within each flank and within the gene body but never across the TSS or polyA site, with the unsmoothed values as thin lines; each of the ten body bins is drawn at the width of one flank bin, so the gene body is not to scale. Values above 15 are clipped (12_figures/06_fig4.py). Per-gene values (Fig. 4f-j) pool the 10 body bins into one maternal and one paternal value per gene, with the same normalisation and averaging over crosses, for genes with at least 10 assigned reads over the body in each cross (all but one UM-biparental gene; 07_compact_profile.py). The panels show maternal minus paternal enrichment per gene (12_figures/06_fig4.py). Within each of the five sets, maternal was compared with paternal by two-sided Wilcoxon signed-rank test. Between sets (gbM-PEG against the demethylated and against all gbM-biparentals, UM-PEG against UM-biparentals), the log2 maternal to paternal ratio, the maternal value alone and the paternal value alone were each compared by two-sided Mann-Whitney U test, so that a maternal gain can be told apart from a paternal loss. All 14 tests were Holm-corrected together (12_figures/06_fig4.py).

### Rice methylation methods

Rice imprinted genes were the persistent imprinted genes of Tonosaki et al., imprinted at 3, 5 and 7 DAP (245 PEGs and 95 MEGs) [30]. Genes were assigned UM, gbM or teM from Nipponbare young leaf methylation (SRR10763664 [39]) with the rules used for maize [18]: UM if mCG and mCHG were both at most 0.05, gbM if mCG was at least 0.2 and mCHG at most 0.05, and teM if both were at least 0.4 (Supplemental Dataset 2). There was no coverage minimum, except that teM also required at least 10 reads over the gene’s CHG sites. The 7 PEGs and 20 MEGs annotated as transposable element related in MSU7 had no leaf methylation values and were left out, leaving 238 PEGs and 75 MEGs. Biparental genes were the genes of each category on none of Tonosaki et al.’s imprinted lists at any stage (their Supplementary Data 3): 8,210 gbM, 16,556 UM and 5,918 teM. Enrichment of gbM among PEGs relative to MEGs was tested by two-sided Fisher’s exact test on the genes with a defined category (gbM against UM and teM).

For allele-resolved methylation, the 5-DAP F1 endosperm whole-genome bisulfite sequencing of Tonosaki et al. (DRA015356) was remapped: Nipponbare x Kasalath, Kasalath x Nipponbare, Nipponbare x 93-11 and 93-11 x Nipponbare, two replicates each (DRR425631 to DRR425638). Kasalath and 93-11 SNPs were called from the resequencing runs Tonosaki et al. used (DRR013715 and DRR013716; SRR933668 and SRR933669) with Bowtie2 2.5.4 and bcftools 1.21. Homozygous sites with at least 5 reads and at least 90% alternative reads were kept (1,883,842 and 1,998,737 SNPs), and these positions were set to N in IRGSP-1.0 for each cross. Reads were trimmed with Cutadapt 5.2 using Trim Galore’s default settings, aligned with Bismark 0.24.2 (-N 1 -L 32, as Tonosaki et al.), deduplicated, and assigned to parent with SNPsplit 0.6.0 (--bisulfite). CG methylation was extracted for each allele with bismark_methylation_extractor (--no_overlap) (13_rice/05_map_libraries.sh). Bisulfite conversion, from chloroplast CHH, was 99.52 to 99.60%.

Parent of origin was compared on the same sequence, as for arabidopsis: the Nipponbare allele as maternal (Nipponbare mother) against paternal (Nipponbare father), and the Kasalath and 93-11 alleles the same way, giving four comparisons. For each gene (MSU7 gene body) and comparison, CG methylation was the weighted ratio on each side, with counts summed over replicates. A comparison counted when both sides had at least 10 covered CG sites, and a gene’s maternal and paternal values were the means over its counting comparisons (07_gene_matpat.py). Supplemental Fig. S6e plots per-gene paternal minus maternal methylation. Maternal and paternal were compared by two-sided Wilcoxon signed-rank test, and gene sets by two-sided Mann-Whitney U test. Supplemental Fig. S6a-d shows maternal and paternal metagenes on the Fig. 2a-d bins. Per comparison and bin, counts were summed over replicates and genes, and a comparison entered a bin when both sides had at least 20 covered CG sites. The drawn values are means over comparisons, and shading is a 95% bootstrap interval over genes, 1,000 resamples (08_metagene.py, 10_figS6.py).

### Arabidopsis methylation methods

Imprinted genes were not re-called from reads. Instead, consensus sets were built from the MEG and PEG lists published by three endosperm screens of reciprocal crosses: Pignatta et al. 2014 (103 PEGs and 285 MEGs, the genes called in any of their three pairs of crosses, Col x Ler, Col x Cvi and Ler x Cvi; Figure 1 source data 2 [41]), Gehring et al. 2011 (43 PEGs and 165 MEGs, Col x Ler; Supplemental Table S6 [42]) and Wolff et al. 2011 (27 PEGs and 39 MEGs, Col x Bur; Supplemental Tables S7 and S4 [43]). Identifiers were reduced to AGI locus codes (isoform suffix removed, upper case), and a gene was a consensus PEG or MEG when it was on the corresponding list in at least two of the three studies, giving 36 consensus PEGs and 70 consensus MEGs (11_arabidopsis/02_imprinting_calls.py, 03_consensus_sets.py; Supplemental Dataset 3). Genes were assigned UM, gbM or teM from the Col-0 leaf classification of Bewick et al. 2016 [22], with its CHG- and CHH-methylated categories combined as teM (05_master_table.py). No biparental set based on expression in endosperm was produced, but the remaining 26,665 genes, which were on none of the three screens’ PEG or MEG lists, had 20,771 UM, 4,851 gbM, and 1,043 teM genes (10_biparental_class_counts.py). Enrichment of gbM among PEGs relative to MEGs was tested by two-sided Fisher’s exact test (11_imprinted_class_enrichment.py).

For allele-resolved methylation analysis in hybrid endosperm, reciprocal Col-0 x C24 wild-type endosperm at 7 DAP, three replicates per direction (GSE280598) [44], was used as deposited per-cytosine methylation-called files on TAIR10. The maternal Col-0 allele came from Col-0 x C24 (GSM8601909 to GSM8601911) and the paternal Col-0 allele from C24 x Col-0 (GSM8601912 to GSM8601914), so the same Col-0 sequence is compared as maternal and paternal and parent of origin is not confounded with ecotype. Gene coordinates were from the Ensembl Plants release 57 TAIR10 annotation (11_arabidopsis/07_gene_targets.py). For each gene and allele, methylated and unmethylated reads were summed over the gene body and the three replicates (08_paternal_allele_methylation.py), and genes with at least 10 covered CG sites on each allele were used (12_figures/09_fig5cde.py). Supplemental Fig. S7e plots whole-gene paternal minus maternal CG methylation for consensus gbM-PEGs, gbM-biparentals and UM-biparentals, tested by two-sided Wilcoxon signed-rank test (12_figures/09_fig5cde.py). Supplemental Fig. S7a-d shows CG methylation metagenes of wild-type Col-0 7-DAP endosperm and young leaf (GSE280598, EM-seq, three replicates each, summed) on the Fig. 2a-d bins, a bin drawn when it has at least 20 covered CGs, n being the most genes behind any bin (12_figures/15_fig_arabidopsis_metagene.py).

For the allele-resolved methylation analysis in *demeter* mutant hybrid endosperm, allele-resolved endosperm methylomes of Ibarra et al. 2012 [45] (GSE38935) were used as deposited per-cytosine files. The *dme* library (GSM952440) is endosperm of aborting seeds from *dme-2*/*DME* Col-gl mothers crossed with Ler; the wild-type library (GSM952441) is from a separate wild-type Col-0 x Ler cross. For each gene and allele, CG methylation was the weighted ratio over the gene body, and a gene needed at least 5 covered CG sites on each allele of the pair compared; the threshold is lower than for GSE280598 because these libraries are shallower (09_dme_fie_ methylation.py). Supplemental Fig. S7f plots whole-gene paternal minus maternal methylation within each library, because the two alleles of one library share its depth, conversion and batch. Within each library, maternal and paternal methylation were compared by two-sided Wilcoxon signed-rank test, and wild-type and *dme* by two-sided Wilcoxon signed-rank test on the per-gene differences of genes measured in both libraries (12_figures/09_fig5cde.py). The two alleles in these libraries are different ecotypes, so unlike the GSE280598 analysis above, parent of origin is not separated from ecotype.

## Statistical analysis

All tests were two-sided and used SciPy. Two independent sets were compared by Mann-Whitney U test, paired per-gene values by Wilcoxon signed-rank test (SciPy default handling of zero differences), proportions by Fisher’s exact test, and more than two categories by Kruskal-Wallis test. Effect sizes are medians of per-gene differences with their IQR, and Cliff’s delta where two sets are compared (lib/stats_lib.py). Holm correction was applied within two figure-level families, stated in their sections: the six DMR-distance tests and the 14 allele-resolved H3K27me3 tests. Other p values are uncorrected, each answering a separate question stated in the text. Bootstrap intervals resample genes with replacement, 1,000 times, and report the 2.5th and 97.5th percentiles. Random seeds are fixed in every script that resamples or shuffles.

## ACKNOWLEDGMENTS

This study was supported in part by resources and technical expertise from the Georgia Advanced Computing Resource Center, a partnership between the University of Georgia’s Office of the Vice President for Research and the Office of the Vice President for Information Technology. This study was funded by NSF grant 2343681 to J.I.G.

## CONTRIBUTIONS

C.D., J.I.G., and Y.S. analyzed data. J.I.G. and C.D. wrote the paper. J.I.G. designed and supervised the study.

