## Supplemental Figures for "Maternal repression of imprinted genes in endosperm by gene body demethylation"

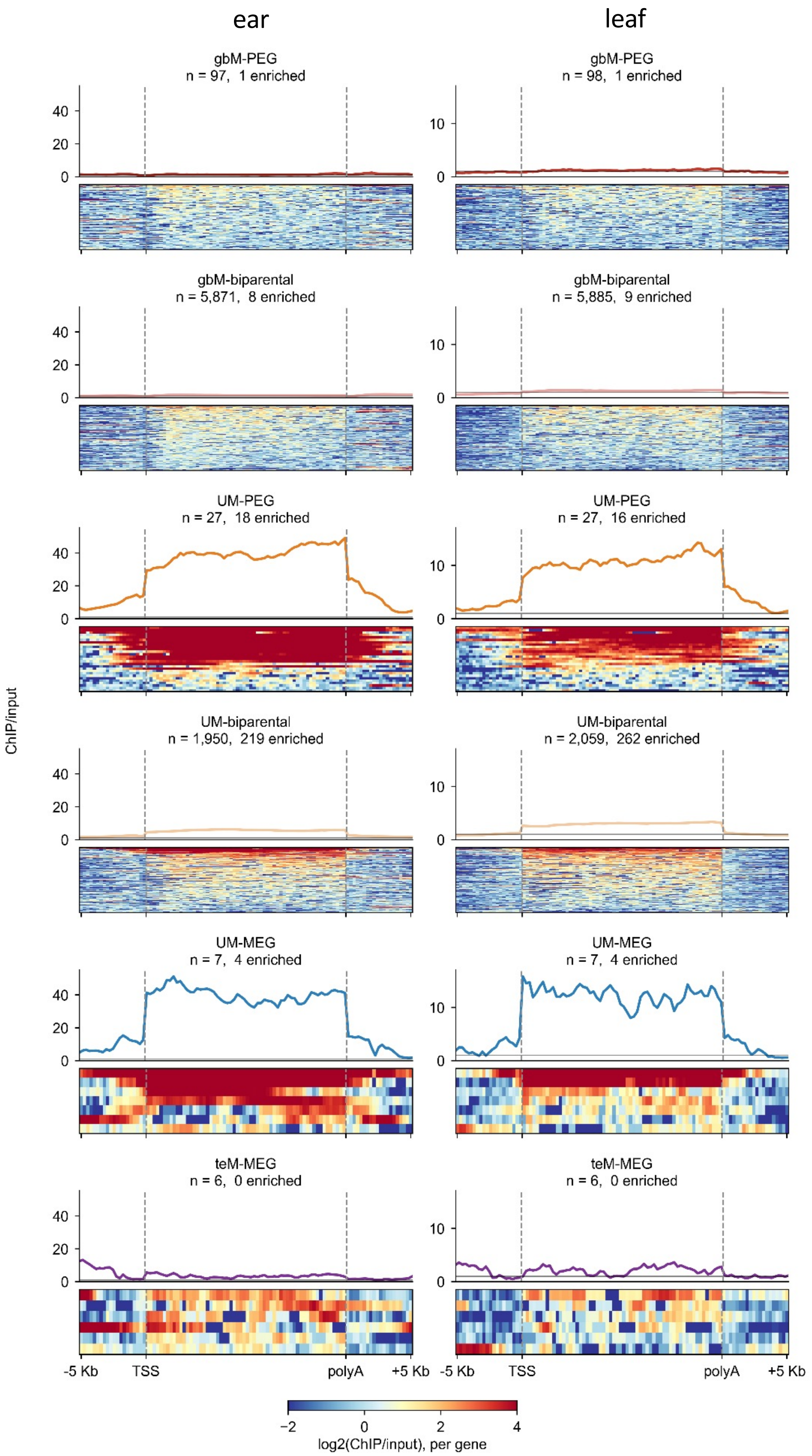


<u>Supplemental Figure S1: Lack of H3K27me3 in gbM-PEGs in leaf and ear</u>

Metagene plots show the normalized ratio of H3K27me3 ChIP-seq reads over input reads. Each gene region was divided into 20 upstream bins of 250 bp each, 60 gene body bins scaled to the gene's length, and 20 downstream bins of 250 bp each. To account for the low number of input reads, enrichment in each bin was measured as ChIP reads per bp in that bin divided by the input reads per bp over the region (flanks included), scaled by the ratio of input to ChIP library sizes. Heat maps show the enrichment values per bin, on a log2 scale. Genes whose ChIP/input ratio over the gene body exceeded a value of 2 were counted as H3K27me3 enriched.

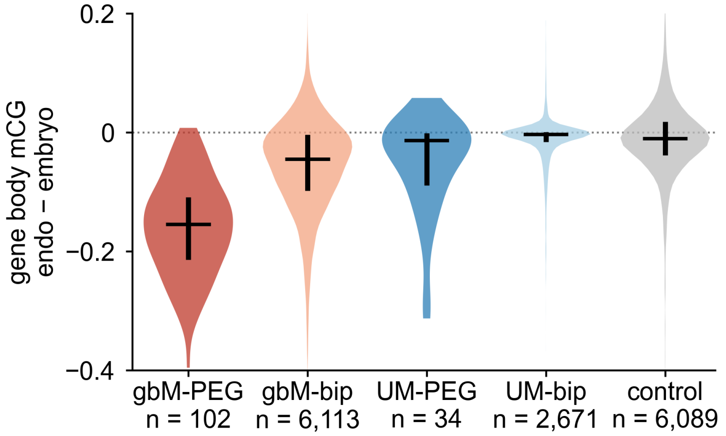


Supplemental Figure S2: Differences in gene body methylation between embryo and endosperm

Violin plots show distribution of CG methylation differences between 15-DAP endosperm and embryo in W22. Methylation was calculated as whole-gene averages. “bip” indicates biparental. Control regions are randomly shuffled from the gbM-biparentals and have the same distribution of region lengths. Horizontal segments indicate median values and vertical segments indicate IQRs. 12 gbM-bip values are outside the y-axis range (11 below -0.4, 1 above 0.2), and 45 control regions are (13 below, 32 above).

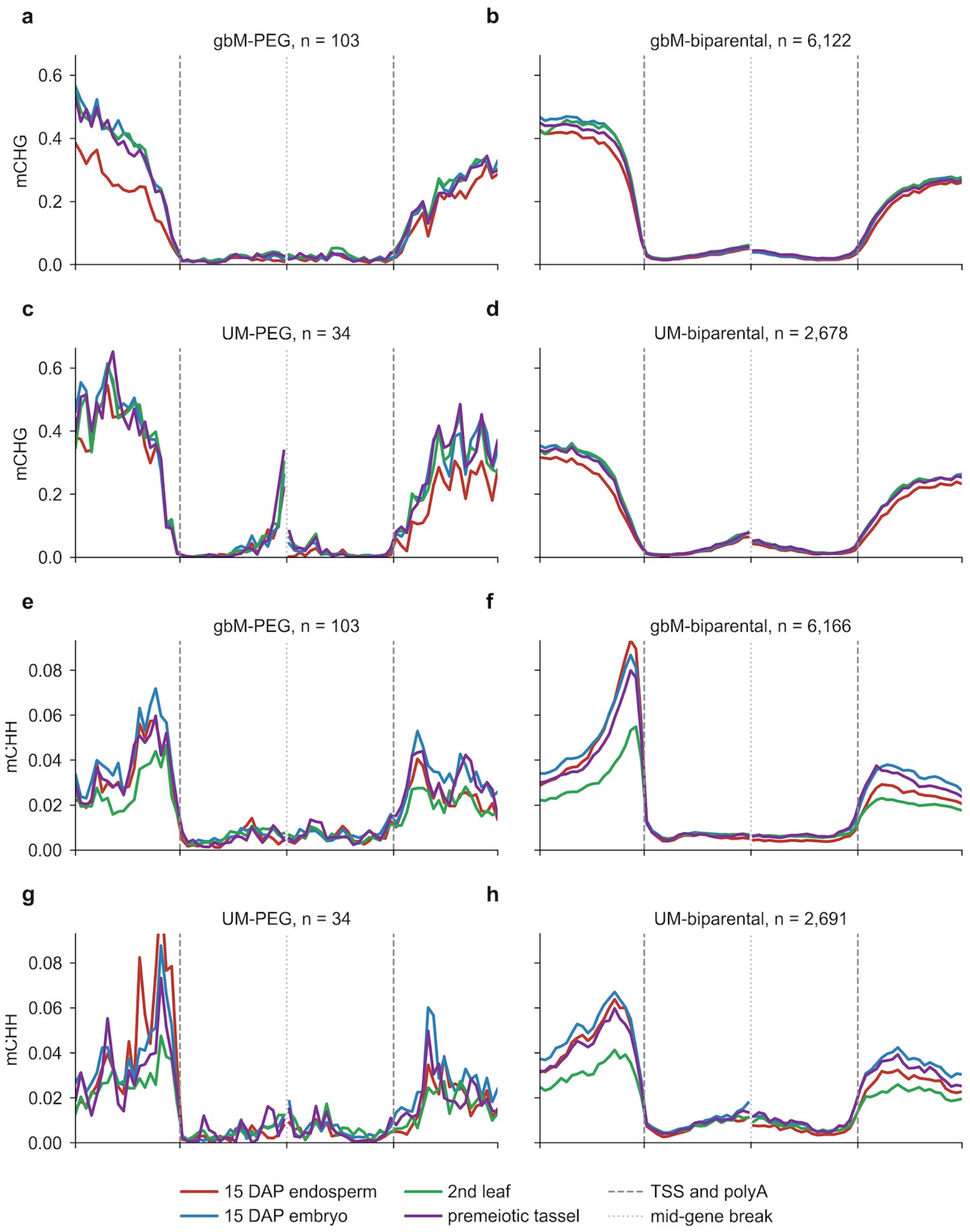


<u>Supplemental Figure S3: CHG and CHH methylation profiles</u>

**a-d.** Metagene plots show methylation trends in wild-type tissues. mCHG is the fraction of methylated cytosines per total cytosines, calculated on 100-bp intervals, 2 Kb internal and 2 Kb flanking genes on each side. Transcription start sites and polyadenylation sites are indicated by vertical dashed lines, mid-gene break by vertical dotted line.

**e-h.** Same as above, but showing CHH methylation.

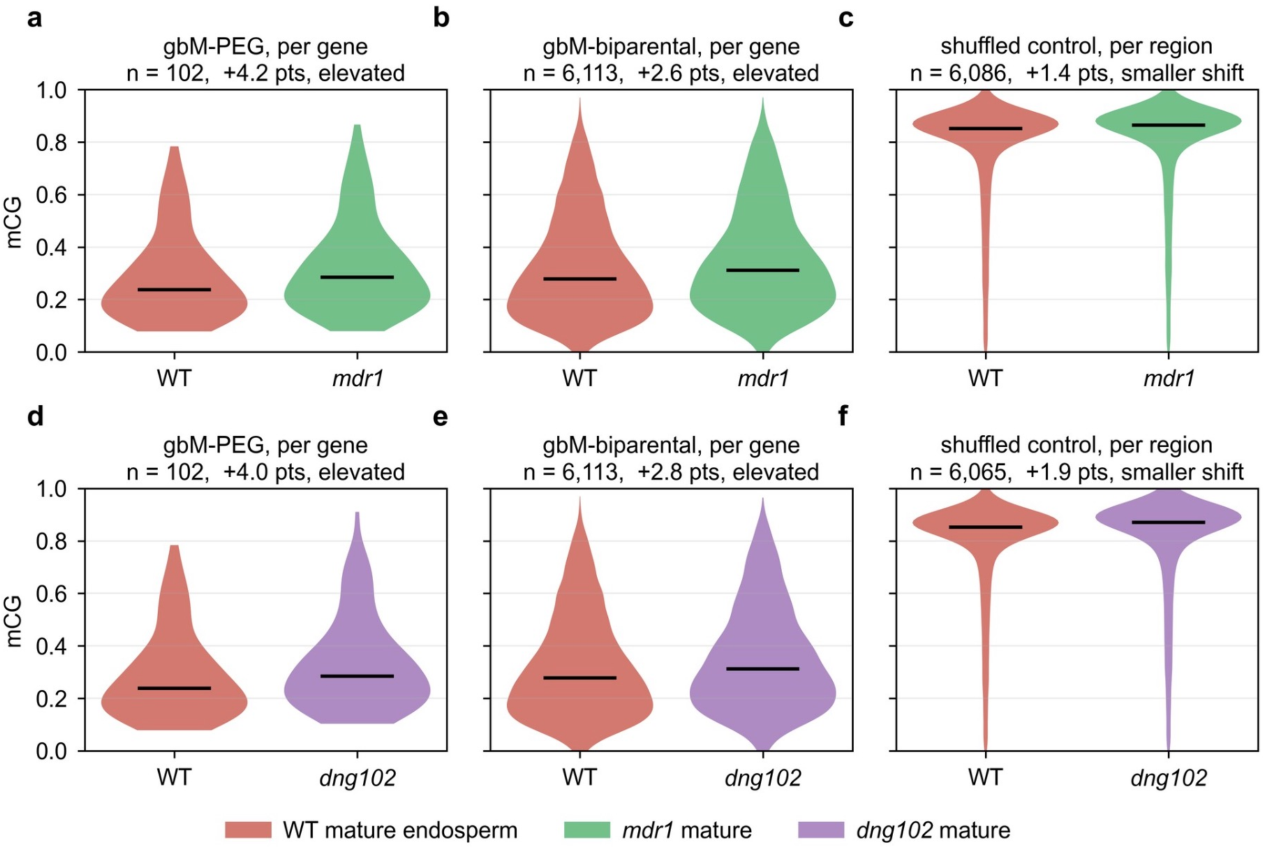


Supplemental Figure S4: Elevated methylation in *mdr1* and *dng102* single mutants in mature endosperm
Violin plots show distributions of CG methylation calculated as whole-gene averages. Horizontal segments indicate medians of the distributions. "pts" is the median of the per-gene differences (per region for the control), mutant minus WT, in percentage points.

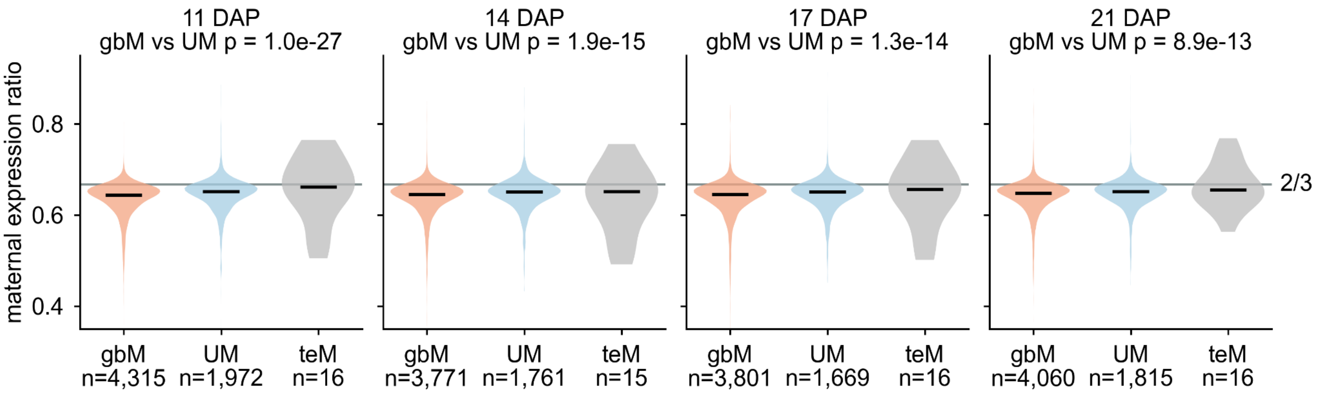


Supplemental Figure S5: Maternal expression ratios based on methylation category

Distributions of maternal expression ratios from B73/W22 hybrid endosperm from bidirectional crosses at four timepoints in endosperm development, showing biparental genes in each methylation category whose maternal plus paternal expression was at least 5 RPM in each direction of the cross, for one-to-one B73/W22 genes, with the ratio averaged over the two directions. Horizontal segments indicate median values. P values are two-sided Mann-Whitney, gbM vs UM.

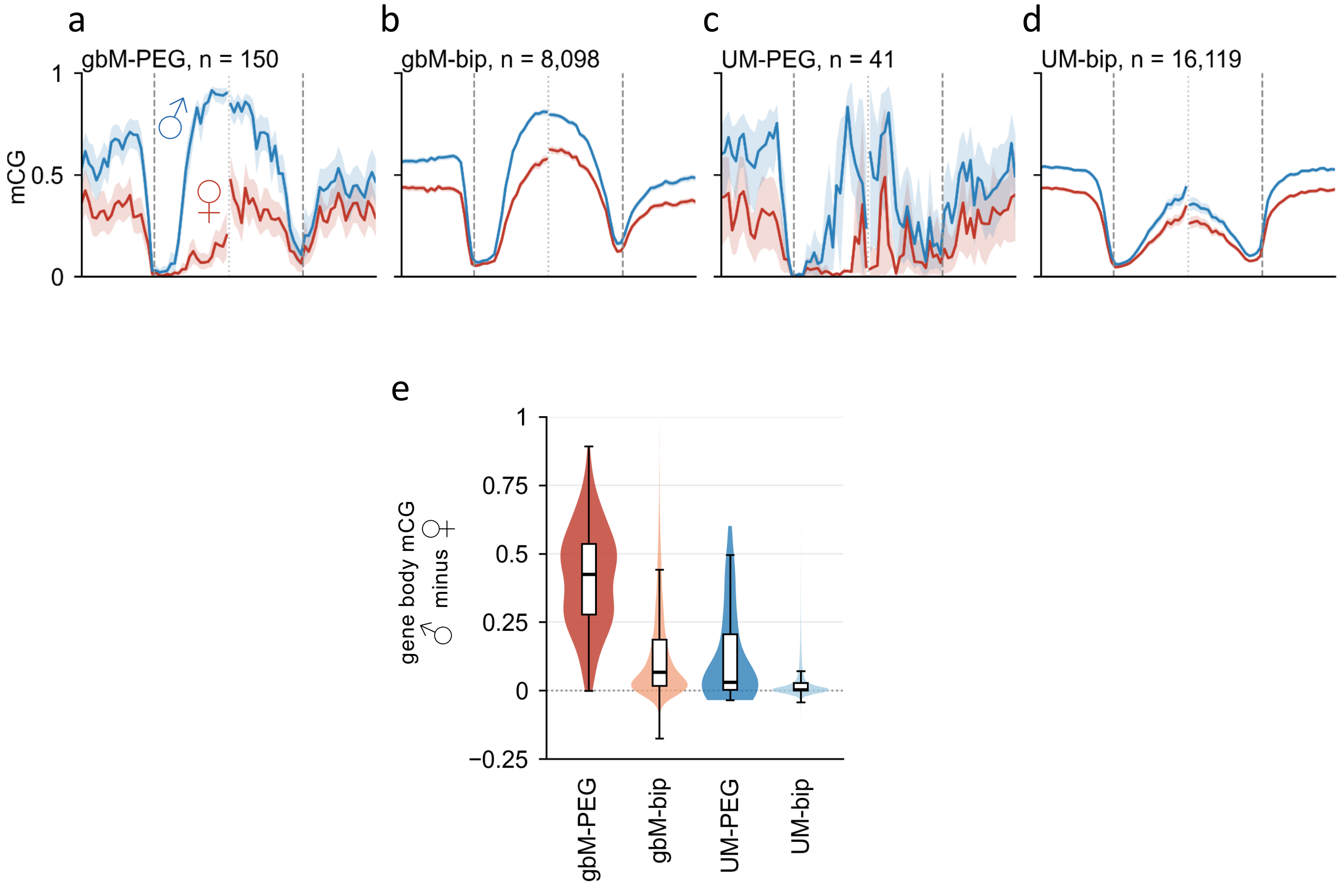


Supplemental Figure S6: Maternal gene body demethylation in rice

**a-d.** Metagene plots show maternal and paternal methylation trends in gbM and UM genes in 5-DAP endosperm. mCG is the fraction of methylated cytosines per total cytosines, calculated on 100-bp intervals, 2 Kb internal and 2 Kb flanking genes on each side. “bip” indicates biparental. Transcription start sites and polyadenylation sites are indicated by vertical dashed lines, mid-gene break by vertical dotted line. Shading indicates 95% bootstrap intervals.
**e.** Parentally resolved methylation shows gbM-PEGs and gbM-biparentals have reduced maternal CG methylation in 5-DAP endosperm from reciprocal crosses of Nipponbare with Kasalath and with 93-11. mCG was calculated as whole-gene averages. The plot shows the distribution of per-gene differences between paternal and maternal methylation, comparing each allele with itself across the reciprocal crosses. Boxes indicate medians and IQRs, and whiskers extend to 1.5 x IQR. 150 gbM-PEGs, 7,922 gbM-bips, 41 UM-PEGs and 15,325 UM-bips with at least 10 covered CGs on both sides of each comparison were included in this analysis.

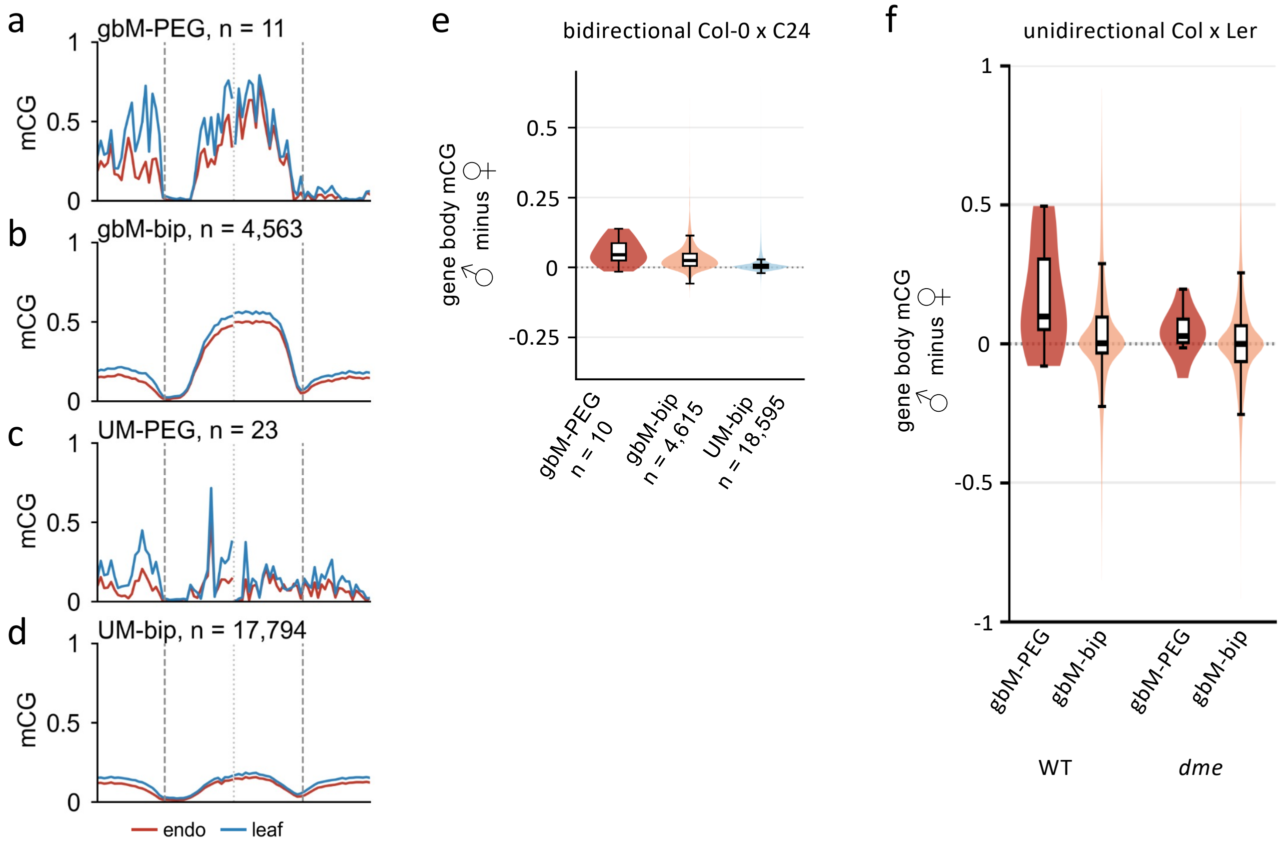


Supplemental Figure S7: Maternal gene body demethylation in *Arabidopsis thaliana*

**a-d.** Metagene plots show CG methylation in wild-type Col-0 7-DAP endosperm and young leaf. mCG is the fraction of methylated cytosines per total cytosines, calculated on 100-bp intervals, 2 Kb internal and 2 Kb flanking genes on each side. "bip" indicates biparental. Transcription start sites and polyadenylation sites are indicated by vertical dashed lines, mid-gene break by vertical dotted line.

**e.** Parentally resolved methylation shows gbM-PEGs and gbM-biparentals have reduced maternal CG methylation in 7-DAP endosperm. Methylome data from Col-0/C24 hybrid endosperm from bidirectional crosses. mCG was calculated as whole-gene averages. Whiskers extend 1.5 × IQR.

**f.** Parentally resolved methylation shows maternal methylation is partially restored in *dme* mutant. Methylome data are from Col-0 × Ler cross hybrid endosperm for WT and from Col-gl (*dme* heterozygote) × Ler for *dme*. mCG was calculated as whole-gene averages. Boxes show medians and quartiles, and whiskers extend to 1.5 × IQR. For gbM-PEGs, paternal minus maternal deltas decrease from 9.9 in WT to 2.8 points in *dme*, p = 0.055. WT includes 8 gbM-PEG and 3,649 gbM-bip; *dme* 9 and 3,839.
